# Cell Painting PLUS reveals phenotypic signatures for nephrotoxicity profiling in human RPTEC/TERT1

**DOI:** 10.64898/2026.10.06.756929

**Authors:** Marlene Wedler, Jose M. Muino, Christopher Wolff, Palle Steen Helmke, Gerhard Ecker, Joshua Harrill, Elena von Coburg, Hugo van Kessel, Giulia Callegaro, Bob van de Water, Michael Oelgeschläger, Shu Liu

**Affiliations:** German Centre for the Protection of Laboratory Animals (Bf3R), German Federal Institute for Risk Assessment (BfR), Berlin, Germany; Institute of Biology, Freie Universität Berlin, Berlin, Germany; Screening Unit, Leibniz-Forschungsinstitut für Molekulare Pharmakologie (FMP), Berlin, Germany; University of Vienna, Vienna, Austria, Department of Pharmaceutical Sciences; Corteva Agriscience, Indianapolis, IN, USA; Leiden University, Leiden Academic Centre for Drug Research, Division of Cell Systems and Drug Safety, Leiden, The Netherlands

**Keywords:** nephrotoxicity, Cell Painting PLUS, high-throughput phenotypic profiling, high-content imaging, in vitro screening

## Abstract

Multiplex high-throughput phenotypic profiling offers a cost-effective approach to monitor how chemical perturbations alter cellular morphology, thereby contributing to a mechanistic, human-centred hazard assessment with new approach methodologies in the context of Next Generation Risk Assessment (NGRA). In this study, we employed Cell Painting PLUS (CPP), an expanded high-content imaging assay that adds spatially resolved channels for mitochondria, lysosomes, actin, Golgi, RNA, DNA, and the endoplasmic reticulum, to renal RPTEC/TERT1 cells to evaluate nephrotoxicity. We profiled a set of nephrotoxic agents together with well-annotated reference chemicals encompassing diverse cellular targets to identify specific morphological profiles associated with distinct mechanisms of action (MoAs). Benchmark concentration modelling, correlation analysis, and hierarchical clustering of RPTEC/TERT1 CPP data captured specific morphological signatures of more than 300 tested compounds and enabled discrimination between closely related and opposing phenotypes. Comparison with CPP data generated in bone osteosarcoma U2OS cells highlighted cell-type-specific differences: despite lower overall activity, RPTEC/TERT1 resolved biologically relevant nephrotoxicity-associated phenotypes, highlighting the importance of using physiologically relevant cell models for context-specific phenotypic profiling. Overall, these findings establish RPTEC-based CPP as a valuable phenotypic profiling platform supporting mechanistic nephrotoxicity assessment for integration into NGRA.

## Introduction

Next Generation Risk Assessment (NGRA) aims to use more human-relevant, fast and efficient toxicity testing approaches for chemicals while minimising the need for animal testing. Proposed NGRA frameworks incorporate new approach methodologies (NAMs) including, for example, in silico methods, high-throughput screening as well as 2D and 3D cell culture models (Thomas et al. 2019; Schmeisser et al. 2023; Leist et al. 2026). In the context of high-throughput screenings, high-throughput transcriptomics and high-content imaging are both increasingly used methods to assess the hazard of chemicals (Nyffeler et al. 2022; Way et al. 2022; Bundy et al. 2024).

High-content imaging methods utilize automated microscopy and image analysis to quantify multiple morphological features of cells to extract multiparametric profiles. Cell Painting (CP) is one commonly used assay that uses fluorescence dyes to generate phenotypic profiles of cells (Bray et al. 2016; Cimini et al. 2023). High-throughput phenotypic profiling (HTPP) using CP has been applied as a powerful tool for a wide range of different purposes in basic research, drug discovery, and toxicology. Examples include the screening of small molecules to identify phenotypic similarities and underlying mechanisms of action (MoAs) (Gustafsdottir et al. 2013; Bray et al. 2016), genetic perturbations to examine gene functions and molecular mechanisms (Chandrasekaran et al. 2025) or industrial chemicals and pharmaceutical compounds for identification of potential hazards to human health and the environment (Nyffeler et al. 2023).

We recently expanded the multiplexing capacity of the CP method by introducing Cell Painting PLUS (CPP), a modified version of the original CP protocol. CPP extends the original CP protocol by adding lysosome staining and spatially resolved imaging channels for actin cytoskeleton, Golgi apparatus, RNA, and endoplasmic reticulum (ER), thereby aiming to improve organelle specificity and phenotypic resolution (von Coburg et al. 2025; Wedler et al. 2026a).

Many CP-based HTPP studies have traditionally been applied to U2OS cells (Gustafsdottir et al. 2013; Bray et al. 2016; Nyffeler et al. 2020; Pahl et al. 2023) thus limiting the use of these data to study specific target organ toxicity, such as nephrotoxicity. Nephrotoxicity is among the major safety concerns leading to rejection of drug candidates during development (Cook et al. 2014; Craveiro et al. 2020). Because tubular epithelial injury is a central mechanism of drug-induced nephrotoxicity, (Kwiatkowska et al. 2021) renal tubular epithelial cell models are particularly relevant for these studies. The human TERT1-immortalized primary renal proximal tubule epithelial cells RPTEC/TERT1 (RPTEC) have proved useful for elucidating mechanisms of drug-induced nephrotoxicity (Simon-Friedt et al. 2015; Pearson et al. 2023), for example, using gene expression analysis (van Kessel et al. 2025) and high-content imaging screens (Ramm et al. 2019). Furthermore, RPTEC has been suggested to be integrated into a NAM-based multi-omics approach for pathway toxicity (Jochum et al. 2025) in support of its value for NGRA approaches.

In the present study, we evaluated the nephrotoxic prediction performance in an established RPTEC CPP assay based on activity and similarity of compounds with shared MoAs using nephrotoxic compounds, cytokines and growth factors as well as a compound set comprising over 300 target-annotated reference chemicals that cover diverse signalling pathways and molecular functions. CPP captured detailed morphological signatures that enabled discrimination between similar and opposing MoAs. Furthermore, we compared the morphological profiles of RPTEC to U2OS cells to investigate cell-line-specific response. Our findings highlight that sensitivity should not be the sole criterion for choosing optimal cell lines for use in the CPP assay. Although RPTEC exhibited lower overall activity than U2OS cells, their physiologically relevant proximal tubule phenotype enabled biologically meaningful characterisation of nephrotoxic mechanisms. Collectively, these findings establish CPP applied to RPTEC as a valuable platform for phenotypic nephrotoxicity assessment and support its application in high-throughput early nephrotoxicity prediction.

## Materials and methods

### Cell culture and cell seeding

U2OS and RPTEC/TERT1 (RPTEC) cells were cultured as previously described (Wedler et al. 2026a): The U2OS (HTB-96) cells were obtained from ATCC and RPTEC cell line was a gift from Bob van de Water (Leiden University, The Netherlands) (van Kessel et al. 2025). The cells were routinely cultured at 37°C and 5% CO2, and absence of mycoplasma was confirmed.

RPTEC were cultured in DMEM (no glucose, sodium pyruvate, no glutamine) (Gibco/Thermo Fisher Scientific, No.11966025) with 50% (v/v) Ham’s F-12 Nutrient Mix (Gibco/Thermo Fisher Scientific, No. 21765029), 2 mM GlutaMAX (Gibco/Thermo Fisher Scientific, No.35050-061), 5 µg/ml insulin, 5 µg/ml transferrin, 5 ng/ml sodium selenite (Sigma-Aldrich/Merck, No.I1884-1VL), 0.5% (v/v) FBS (Tico Europe, No.FBSEU500), 10 ng/ml hEGF (Sigma-Aldrich/Merck, No.E9644-2MG), 36 ng/ml hydrocortisone (Sigma-Aldrich/Merck, No. H0135-1MG), 100 µg/ml streptomycin / 100 U/ml penicillin (PanBiotech, No. P06-07100), and 0.1 mg/ml geneticin (G418) (Roche, No. 04727894001).

U2OS culture medium was composed of DMEM (4.5 g/l glucose, sodium pyruvate, glutamine, no phenol red) (PanBiotech, No. P04-03588) supplemented with 5% (v/v) FBS (Sigma, No. BCBQ7890V) and 100 µg/ml streptomycin / 100 U/ml penicillin (PanBiotech, No. P06-07100).

For RPTEC, 384-well Screenstar plates (Greiner Bio-One) were pre-coated with collagen (0.1 mg/ml collagen type I isolated from rat tail, Ibid, #50201) in water for 1 h, then washed once with 40 µl PBS and seeded 6,000 cells/well in 40 µl/well. RPTEC were grown 10 d on plates post-confluence to enable contact inhibition–driven maturation, resulting in a polarized epithelial monolayer with characteristic cobblestone morphology (Wieser et al. 2008).

Cells were passaged 2-3 times before being seeded onto plates. For U2OS, 3,000 cells/well in 40 µl/well were seeded in 384-well PhenoPlates (Revvity Inc.) and grown for 24 h at 37°C and 5% CO_2_ before compound treatment.

### Compound selection

Nephrotoxic compounds, growth factors and cytokines were selected based on a previous publication investigating mechanistic assessment of nephrotoxicity *in vitro* in RPTEC (van Kessel et al. 2025). Additionally, a curated set of 310 bioactive compounds with annotated molecular targets was assembled from RefChemDB as part of a Tox21 consortium cross-partner project between the U.S. Environmental Protection Agency (EPA) and the National Institute of Environmental Health Sciences (NIEHS DTT) (Tox21 Consortium https://tox21.gov/projects/; Judson et al. 2019) as described in (van Kessel et al. 2026). Subsequently, this Tox21 compound set will be referred to as ‘reference chemicals’. Where possible, a minimal set of two compounds per target protein was selected to enable comparative analysis. Because compound function depends on both the target protein and the mode of modulation, target identity and modulation (agonist or antagonist) were jointly defined as the compound MoA (Supplementary Table 1). Following quality control by mass spectrometry, 26 compounds were excluded because the expected molecular mass was not detected (van Kessel et al. 2026). The CPP screen data for the remaining 284 compounds were used for further analysis. Furthermore, a set of 15 CPP performance control compounds was included, containing positive and negative controls for CPP (von Coburg et al. 2025).

### Compound treatment

The annotated reference chemical set was obtained from the US EPA’s chemical contractor EvoTec (Princeton, NJ, USA) and supplied as stock solutions in Labcyte Echo-qualified 384-well low-dead-volume (384LDV) plates. Stock solutions were prepared at 20 mM or, for compounds with limited solubility, at the highest concentration achievable in DMSO (maximum 20 mM). Seven-point half-log serial dilution series were prepared for compound testing. Nephrotoxic compounds, growth factors, and cytokines were prepared as previously described in seven-point dilution series (van Kessel et al. 2025). CPP performance control compounds were prepared as previously described (von Coburg et al. 2025). Assay-ready plates were generated using an Echo 650 acoustic liquid handler (Beckman Coulter, Brea, CA, USA).

For the U2OS screen, most compounds were transferred directly from the stock solutions using a 667-fold dilution with a Janus^TM^ liquid handler (Revvity Inc.) to achieve the desired final test concentrations. For some compounds, lower concentrations than those used in the RPTEC screen were used in U2OS cells to avoid cytotoxicity. A detailed description of the U2OS screen is provided in the companion paper (Wedler et al. 2026b, in preparation). Wells containing 0.15% (v/v) DMSO served as vehicle controls. Cells were exposed to the compounds for 24 h at 37°C in a humidified atmosphere containing 5% CO_2_.

The CPP screens of the annotated reference chemical set in RPTEC and U2OS that were analyzed in this study have also been reported in companion studies focusing on different analyses (van Kessel et al. 2026) (Wedler et al. 2026b, in preparation).

### Cell Painting PLUS

Cell Painting PLUS was performed as previously described in detail (von Coburg et al. 2025; von Coburg and Dunst 2026; Wedler et al. 2026a). In brief, CPP comprises a first staining and imaging cycle and a subsequent elution step followed by a second staining cycle. For the first staining cycle, media was aspirated from the wells, leaving only ∼15 µl of residual volume. Then, 15 µl of 2× live cell staining solution with MitoTracker Orange CMTMROS (1.0 μM in staining solution; 0.5 μM in well) and lysosome staining (LysoBrite NIR Fluorescence, 1:250 in staining solution buffer A, 1:500 in well) with LysoBrite NIR Fluorescence buffer B containing 1% BSA were added and incubated for 30 min at 37°C. To fix the cells, 30 µl of 8% PFA in PBS was directly added (resulting in 4% PFA in the wells). After 20 min incubation at ambient temperature, protected from light, plates were washed three times with 100 µl PBS, including final aspiration to leave ∼15 µl PBS inside the wells. To permeabilise and simultaneously stain cells with fixed-cell dyes, 15 µl of 2× staining solution with Triton X-100 (0.2% Triton in staining solution; 0.1% in well), actin stain Alexa Fluor Plus 405 Phalloidin (66.0 nM in staining solution; 33.0 nM in well), and RNA stain SYTO 14 Green Fluorescent Nucleic Acid Stain (0.6 μM in staining solution; 0.3 μM in well) (in PBS containing 1% BSA; resulting in 0.5% BSA in well) was added. After 30 min, plates were washed three times with 100 µl PBS. Subsequently, plates were imaged. After imaging of the first staining cycle, plates were washed three times with 100 µl water before adding 40 µl of elution buffer (composed of 0.5 M L-glycine and 1% SDS in water with pH adjusted to 2.5). After 10 min incubation, plates were washed three times with 100 µl water and three times with 100 µl PBS. Then, the wells were aspirated to a residual volume of ∼15 µl, and 15 µl of 2× staining solution containing the dyes of the second staining cycle was added: Hoechst 33342 (8.0 μM in staining solution; 4.0 μM in well), Wheat Germ Agglutinin Alexa Fluor 488 Conjugate (2.5 μg/ml in staining solution; 1.3 μg/ml in well), and Concanavalin A Alexa Fluor 647 Conjugate (20.0 μg/ml in staining solution; 10.0 μg/ml in well) (in PBS containing 1% BSA; 0.5% in well). After incubation for 30 min at room temperature, the plates were washed three times with 100 µl PBS and imaged.

### Imaging

Plates were sequentially imaged with an Opera Phenix High-Content Screening System (Revvity Inc.) in confocal mode with 2×2 pixel binning using a 20x water objective (NA 1.0) and 4 excitation wavelengths as described previously (von Coburg et al. 2025; Wedler et al. 2026a). For RPTEC and U2OS, 4 and 6 fields/well were imaged, respectively.

### Image Analysis

CPP images were analysed using customised Analysis sequences in Harmony software v4.8 (Revvity Inc.) as described previously in detail (von Coburg et al. 2025; von Coburg and Dunst 2026; Wedler et al. 2026a) with one optical section (z-plane) employed for image analysis. In short, illumination of images was corrected using the flatfield correction option integrated in Harmony. Then, for CPP, single multi-stack images combining images from the first and second staining cycles were created using the application-specific building block *Add Channels 4i* in Harmony (Kirsch 2025). Multi-stack images were then fed in the customized and publicly available Harmony image analysis sequence (von Coburg et al. 2025; Wedler et al. 2026a): In brief, cell segmentation is conducted by first identifying nuclei based on Hoechst staining. Then, cell outlines are detected using merged mitochondria and RNA channel, followed by definition of the cell regions (cytoplasm, membrane, ringregion and nucleoli). Then, feature modules (e.g., morphology, texture, intensity, shape and position) were extracted for each meaningful combination of cell region and channel resulting in 894 morphological features at the single-cell level.

### Data analysis

All data analysis in this study was performed using R [v4.2.2] (R Core Team 2024), Jupyter Notebook (Kluyver et al. 2016) and KNIME [v5.4] (Berthold et al. 2007). The *CPPAnalyzer* Jupyter notebook for data normalisation and the *CPPManager* KNIME workflow for downstream visualisation and BMC calculation have been published previously (Wedler et al. 2026a). R scripts and KNIME workflows used for this study are publicly available (see supplementary information).

### Data normalization

All data were normalised using the published *CPPAnalyzer* Jupyter notebook (Wedler et al. 2026a). For RPTEC CPP data, plate effects were corrected using B-score normalisation as described previously (Mpindi et al. 2015) and implemented for CPP (Wedler et al. 2026a): In brief, the *loess* function from the R package *loess* is applied to well-level aggregated features to remove spatial row-and column plate effects. Then, the data were normalised to the median and median absolute deviation (MAD) of the solvent-control DMSO wells from the plate. The resulting normalised feature values are termed as ‘B-score’ in the following.

For U2OS, nuclei sizes were filtered using a *nucleus area* range of 120-1000 µm^2^ to exclude dead cells or cells with fragmented nuclei. For the U2OS CPP data, no obvious plate effects were observed, and therefore, standard well-level normalisation was applied for all U2OS data as described in (Wedler et al. 2026a). In brief, aggregated well-level data are normalised to the median and MAD of the solvent-control DMSO wells (Bray et al. 2016; Cimini et al. 2023). The resulting normalised feature values are termed ‘robust z-score’ in the following.

For further data cleaning, features with ≥ 15% missing values were excluded, e.g., the features (Cells - Morphology_Nucleus Width [µm]), (Nucleoli - Morphology_Nucleolus Area [µm^2^]), (Nucleoli - Morphology_Nucleolus Width [µm]), (Nucleoli - Morphology_Nucleolus Length [µm]) and (Nucleoli - Morphology_Nucleolus Ratio Width to Length) were excluded for U2OS. Therefore, the final feature lists for RPTEC and U2OS contained slightly different features. Next, objects (single cells) still containing missing values in the remaining features were excluded from the analysis. For downstream analysis, cytotoxic conditions were excluded if the percentage of total cells relative to the DMSO control was below 50% in more than half of the biological replicates.

### Identification of active conditions

Active conditions were identified using the normalised data (robust z-scores or B-scores) calculated as the median across biological replicates. Features were considered active if the absolute robust z-score or B-score was above or equal to 3, indicating a deviation greater than three times the median absolute deviation (MAD) from the median value of the DMSO controls. A treatment condition (compound at a given concentration) was classified as active if at least 1% of features were active.

### Percent replicating

The percent replicating score, used as a reproducibility metric, was defined as previously described with modifications (Way et al. 2022; Wolff et al. 2025). Briefly, for each treatment condition (compound and concentration), pairwise Pearson correlations across all replicates were computed and aggregated using the median. A null distribution was generated from 10,000 randomly sampled profile pairs from the full dataset. Profile pairs belonging to the same chemical were excluded from this sampling. The percent replicating score was defined as the proportion of profile pairs for the same condition whose median Pearson correlation exceeded the 0.95 quantile of the null distribution. Only non-cytotoxic concentrations were used for the percent replicating score.

### Morphological profile heatmaps

Normalised feature values (B-scores or robust z-scores) are used to calculate the median across all biological replicates of a tested condition (compound at a given concentration). Single features are ordered from left to right according to imaging channel, region and module. Normalised feature values are plotted in a heatmap where the colour encodes the B-score or z-score of the respective feature. For visualisation, values are clipped to the range [-6, +6], with values greater than 6 are set to 6 and values smaller than −6 are set to −6.

### Benchmark concentration (BMC) modelling

For the RPTEC CPP screen, seven concentrations were tested, enabling benchmark concentration (BMC) modelling for each feature and compound, as previously described (von Coburg et al. 2025) using a customised KNIME workflow (Wedler et al. 2026a). Briefly, five curve-fitting models (*cnst, hill, poly1, poly2, pow*) from the R package *tpclfit2* (Sheffield et al. 2022) were applied, with the benchmark response (BMR) defined as 1.349*SD. The best-fitting model was selected for each feature. BMCs were defined as inactive if the hitcall was below 0.9, if BMDU or BMDL values were missing, or if the BMC exceeded the maximum non-cytotoxic concentrations.

BMC results were visualised using a BMC proportion map (at the channel level) and an accumulation plot (feature category level). In the BMC proportion map, the proportion of features within each feature category showing significant responses (p < 0.05, one-sided Student’s t-test) to a compound compared with all tested compounds was displayed. For the accumulation plots, only feature categories with at least 30% active BMCs were included. Each data point in the accumulation plot represents a feature category plotted by its ranked median category BMC as described for CPP in (von Coburg et al. 2025; Wedler et al. 2026a).

### Evaluation of Target expression status

mRNA expression data for RPTEC and U2OS were extracted from the Human Protein Atlas (https://www.proteinatlas.org) (Uhlen et al. 2015; Jin et al. 2023). A respective target of a compound was considered as not expressed if the transcripts per million (TPM) value was zero. Note that for Fig. 4 and 6, only compounds whose targets are expressed were considered.

### Comparison between cell lines

For the EPA compounds screened in RPTEC and U2OS, only the concentration tested in both cell lines was considered for comparison. Cell, nucleus size and neighbor contact area were extracted based on the features “Cells - Morphology_Cell Area [µm^2^])”; “Cells - Morphology_Nucleus Area [µm^2^]” and “Cells - Position_Cell_NeighborContactArea Contact Area with Neighbors [%])” from the Harmony CPP image analysis sequences described above. Activity (% active features) and cytotoxicity were determined as described above.

### Mechanism of action (MoA) analysis – Correlation scatter plot

To evaluate correlations between compounds sharing the same MoA, median Pearson correlations between morphological profiles of the compounds sharing the same MoA were calculated and plotted in Fig. 6a. For this purpose, only targets expressed in both U2OS and RPTEC, as described above, were retained for analysis. Based on RefChemDB (Judson et al. 2019), several compounds target multiple proteins, this was included in the Pearson correlation-based MoA analysis for potential polypharmacology.

## Results

### Cell Painting PLUS (CPP) profiling in RPTEC/TERT1: screening overview and quality assessment

To evaluate the predictive performance of RPTEC using the CPP assay for nephrotoxicity, a set of 42 nephrotoxicants, including well-characterised nephrotoxic compounds, cytokines and growth factors was selected (van Kessel et al. 2025). These nephrotoxicants act through different mechanisms, including, e.g., DNA damage, ER stress or oxidative stress.

Furthermore, an annotated reference chemical set was assembled based on RefChemDB (a database from the U.S. EPA) as described in (van Kessel et al. 2026): This reference chemical set comprises a curated set of 310 bioactive compounds with narrowly defined MoA comprising molecular targets and mode of modulation (antagonist/agonist). Where possible, two compounds per target protein were selected to enable comparative analysis. For compounds annotated with multiple targets, the target with the highest support level, defined as the best-characterised target with the highest certainty of relevance, was retained (column “MoA_curated” in Table S1) (Judson et al. 2019). Following quality control by mass spectrometry described in our accompanying study, 26 compounds were excluded because the expected molecular mass was not detected (column “MS_Identified” in Table S1) (van Kessel et al. 2026). The remaining 284 reference chemicals were used for final morphological analysis in RPTEC. In addition, 15 performance control compounds for CPP (von Coburg et al. 2025) were included. The test compounds were screened in RPTEC in a concentration-response format using half-log dilution steps (Fig. 1a). The tested concentration ranges of nephrotoxicants and reference chemicals in the RPTEC CPP screen were selected based on cell viability data for RPTEC described in (van Kessel et al. 2025) and our accompanying study (van Kessel et al. 2026).

**Fig. 1.**
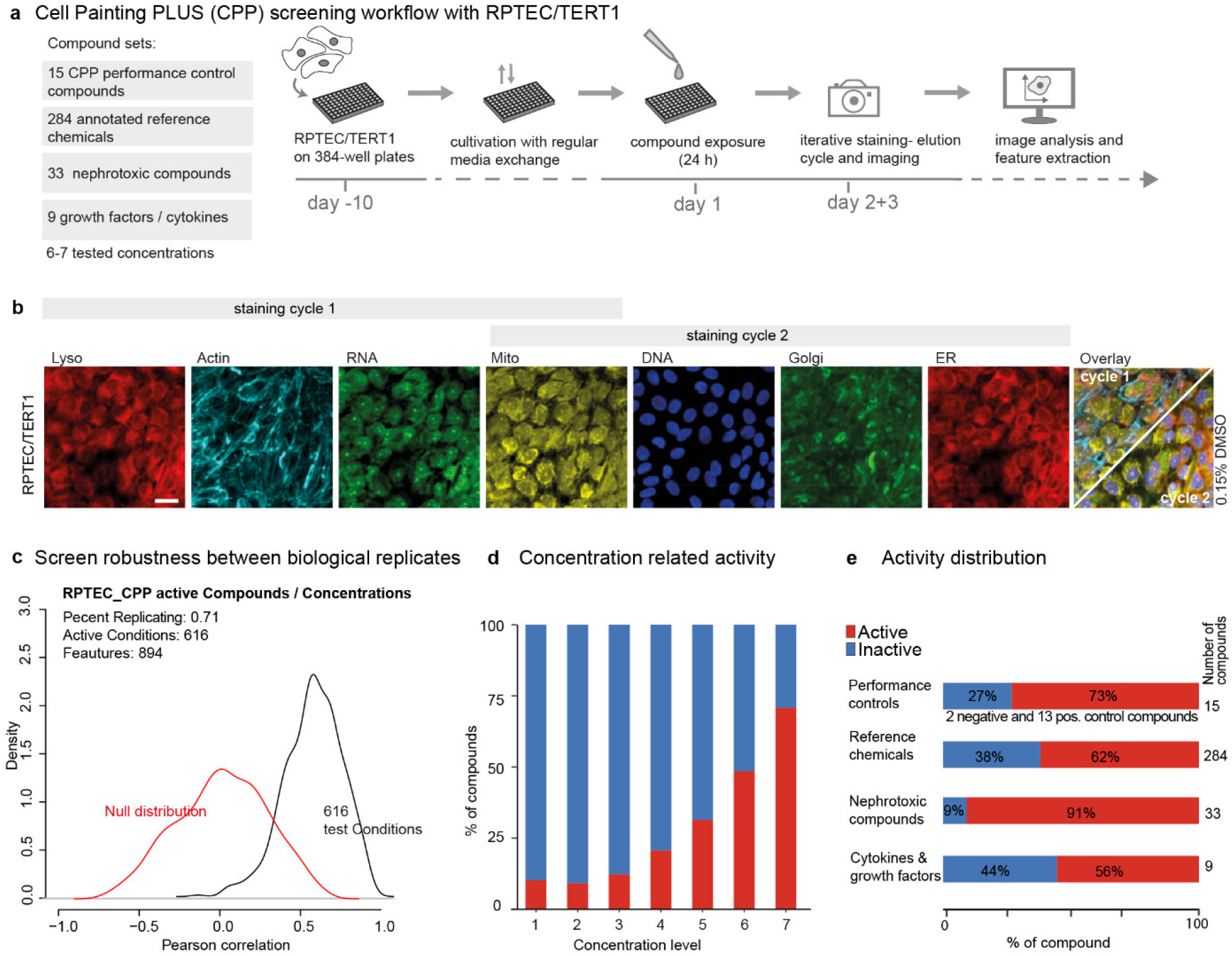
Overview of the Cell Painting PLUS screen in RPTEC/TERT1. **a** Workflow of the Cell Painting PLUS (CPP) screen with RPTEC cells. Cells were seeded in multi-well plates, cultured for 10 days for differentiation and then exposed to test compounds for 24 h. Compounds were screened in concentration-response series in 5 independent biological replicates (n = 5). Cells were subjected to an iterative staining-elution cycle for CPP staining and imaged. From the obtained images, morphological features were extracted. **b** Representative images of RPTEC cells in solvent control conditions (0.15% DMSO) showing the dyes used in the iterative staining cycles of CPP: lysosomes (Lyso), RNA and actin cytoskeleton are stained in cycle 1; DNA, Golgi apparatus and endoplasmic reticulum (ER) in cycle 2. Mitochondrial (Mito) staining is present in both staining cycles to facilitate image registration. The mitochondria-related features are extracted from cycle 2 images. Scale bar = 20 µm. **c** Percent replicating of active compounds at non-cytotoxic concentrations across all five biological replicates. A compound was classified as active when ≥1 % of features displayed a B-score ≥ 3 (representing responses ≥ 3×MAD of the DMSO control). **d** Proportion of active non-cytotoxic compounds across the tested concentrations (level 1 = lowest tested concentration, level 7 = highest tested concentration). Activity increased with higher concentrations. **e** Total number of compounds and the fraction of active compounds at the maximum non-cytotoxic concentration for each compound set; the nephrotoxic compounds exhibit the highest fraction of active conditions.

We used RPTEC for CPP as previously described (von Coburg et al. 2025; Wedler et al. 2026a): RPTEC were cultivated for 10 days to allow them to differentiate, before the cells were exposed to test compounds for 24 h followed by CPP staining and imaging (Fig. 1a). The CPP dye panel comprises fluorescence staining for lysosomes, mitochondria, RNA, actin cytoskeleton, DNA, Golgi and endoplasmic reticulum (ER), each imaged in a separate imaging channel (Fig. 1b). This is accomplished through an iterative staining elution cycle: after imaging of dyes from staining cycle 1, the dyes are eluted to allow re-staining with a second set of dyes in the staining cycle 2. Mitochondrial staining is present in both staining cycles to allow image registration from both cycles (von Coburg et al. 2025).

Screen robustness was quantified by the percent-replicating metric (Way et al. 2022; Wolff et al. 2025), which is based on the median pairwise Pearson correlation of (biological) replicate profiles. As low reproducibility is typically observed for concentrations that induce weak responses (Wolff et al. 2025), only non-toxic and active compound/concentrations were utilised. Non-toxic concentrations were defined as relative percentage of total cells to solvent control being ≥ 50% in more than half of the biological replicates. Active compounds (at a given concentration) were identified as those with ≥1 % of features having an absolute robust z-score or B-score ≥ 3 (representing activities ≥ 3×MAD of the DMSO control). Each condition was tested in five biological replicates. Test conditions (identical compound and concentration across biological replicates) were compared against a null distribution generated from randomly paired profiles. RPTEC CPP screens yielded a percent replicating of 0.71, indicating good screening robustness (Fig. 1c).

Screen quality was assessed with diagnostic plots generated from the raw data using the published *CPPAnalyzer* Jupyter notebook (Wedler et al. 2026a). Analysis of nuclear area and DNA intensity histograms across all five biological replicates (40 plates in total) revealed a single homogeneous cell population (Supplementary Fig. S1a). However, pronounced plate-effect patterns were evident in unnormalised raw data, as indicated by a fluorescence intensity gradient from the left to the right side of the screening plates (Supplementary Fig. S1b). To correct for these plate effects, B-score normalisation was applied as described before (Mpindi et al. 2015). Compared with standard well-level normalisation, B-score normalisation improved correction of systematic artefacts (Supplementary Fig. S1b) and yielded higher percent replicating scores for active conditions (0.71 for B-score normalised data (Fig. 1c) versus 0.63 for standard well-level normalised data (Supplementary Fig. S1c)). Importantly, strong plate-specific patterns observed for negative control compounds (saccharin and sorbitol) were effectively reduced by B-score normalisation (Supplementary Fig. S1d). Therefore, B-score normalisation was used for all analyses of the RPTEC CPP data.

As expected, the number of active compounds increased with rising concentrations (Fig. 1d). When examining the activity for the different compound collections at the maximum non-cytotoxic concentration, the nephrotoxic compound set showed the highest activity, with 91% of the nephrotoxic compounds identified as active. Cytokines and growth factors showed the lowest percentage of active compounds (56%), suggesting that the screening conditions may not have been optimal for this compound class (Fig. 1e). Potential explanations include reduced bioavailability of these proteins during the 24 h incubation period due to protein instability or cellular uptake, and a treatment duration that may have exceeded the optimal window for detecting the induced morphological changes, as many cytokines and growth factors elicit rapid and transient signalling responses that peak within minutes to hours (Braun et al. 2013; Yamagishi et al. 2024), e.g., transcriptional response to cytokines and growth factors in RPTEC has been observed within 4 to 8 hours (van Kessel et al. 2025).

The compound sunitinib was included in different compound sets (e.g., in the reference chemicals set and in the performance control compound set) from distinct sources and thus tested twice within the RPTEC CPP screen. Paired comparison exhibited a high Pearson correlation coefficient of 0.84 for sunitinib from the different test batches, confirming the reproducibility of the screen and the concordance between different batches of compounds (Supplementary Fig. S2).

In summary, we demonstrated good robustness for the implemented CPP screen. Analysis of the morphological profiles revealed high activity for nephrotoxic compounds. Next, we aimed to assess the predictive performance for nephrotoxicity using CPP in RPTEC, focusing on the ability of CPP to identify the activity of nephrotoxic compounds across different organelles.

### CPP profiling recapitulates organelle-specific phenotypes of nephrotoxic compounds

First, we examined the activity of nephrotoxic compounds across all tested concentrations. At the lowest tested concentration, almost all compounds showed no activity, while activity increased in a concentration-dependent manner for most compounds (Fig. 2a). Ochratoxin A, tunicamycin, and amphotericin B induced large phenotypic activity already at the second- and third-lowest tested concentrations. In contrast, citrinin, lead acetate, N-phenylanthranilic acid (NPA), and tetracycline elicited notable activity only at the two highest tested concentrations, which may reflect the requirement for higher intracellular concentrations to induce measurable morphological alterations. Cadmium dichloride and potassium dichromate, both well-established nephrotoxic heavy metals, produced the strongest phenotypic responses, with more than 80% active features at the highest tested concentration (MaxConc). Heavy metals are known to induce oxidative stress, DNA damage and mitochondrial dysfunction (Balali-Mood et al. 2021). Conversely, tenofovir, phenacetin and paracetamol did not induce pronounced phenotypic activity at the highest tested concentration (Fig. 2a). This possibly reflects their comparatively weak acute morphological effects under these experimental conditions, as previous studies have shown that toxic effects of phenacetin (Kobayashi et al. 2012; Taxak et al. 2013), paracetamol (Potter and Hinson 1987; Athersuch et al. 2018) and tenofovir (Delaney et al. 2006; Hamlin et al. 2019) may require metabolic activation. Moreover, mitochondrial effects of tenofovir have been reported in RPTEC following 24 h exposure to 300 µM tenofovir (Pearson et al. 2023), which is substantially higher than the maximum concentration tested in the present study (MaxConc, 20 µM).

**Fig. 2.**
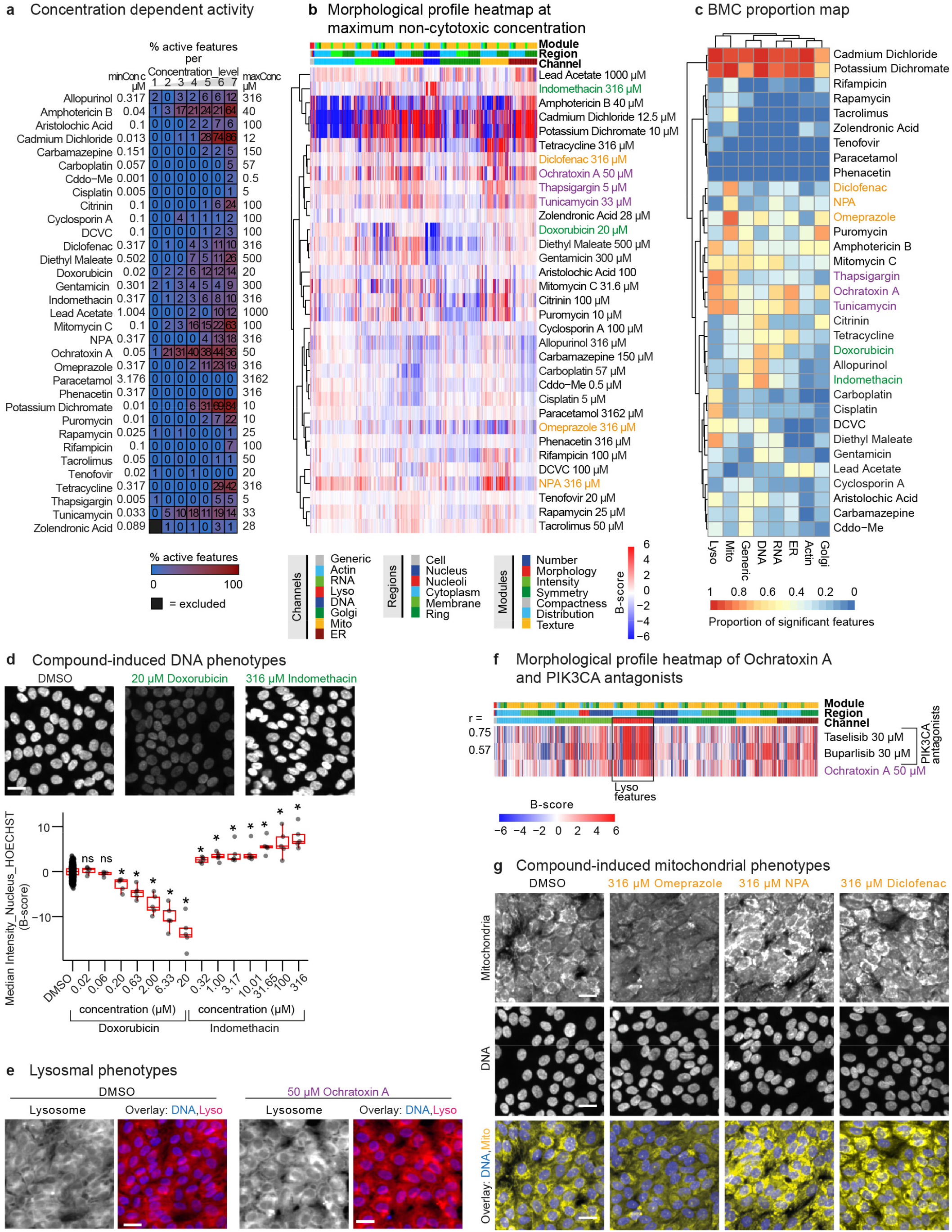
Profiling of nephrotoxic compounds in RPTEC/TERT1. **a** Percentage of active features (absolute B-scores ≥3) for all tested concentrations of the compounds. Each row corresponds to a single compound, columns represent concentration levels 1 (lowest) to 7 (highest). The minimum (minConc) and maximum (maxConc) tested concentrations are given in µM, with serial dilutions performed in half-log steps. Numbers and colour inside the squares indicate the proportion of active features at that concentration (%). Black squares denote conditions excluded due to technical errors. **b** Morphological profile heatmap representing normalised feature values (B-scores) for compounds at their maximum non-cytotoxic test concentration. Median values from 5 biological replicates are shown. Each column represents a single morphological feature; colour encodes the B-score of that feature (clipped to the range [-6, +6], values > 6 are set to 6 and values < −6 are set to −6 for display). Modules, cellular regions and imaging channels are colour-coded in the top legend. The accompanying dendrogram shows hierarchical clustering of compounds based on Pearson correlation of their feature vectors. Colours in compound name labels indicate compounds specifically discussed in the main text. **c** BMC proportion map indicating proportion of single features that show significant response to the specific compound compared to all other compounds (p<0.05) within each channel. Columns are hierarchically clustered using the Ward.D2 linkage method. **d** Representative images from the DNA channel (Hoechst-stained) for RPTEC cells treated with doxorubicin, indomethacin and DMSO solvent control. Scale bar = 20 µm. Box plots illustrate concentration-dependent changes in nuclear intensity; individual points represent single-well measurements pooled from all replicates (n = 5). Asterisks indicate significant differences from DMSO (Kruskal-Wallis test with Dunn’s post-hoc test with Bonferroni correction; *p < 0.05; ns = not significant). **e** Representative images of ochratoxin A-treated cells and DMSO solvent control: lysosome and DNA stains are displayed. Scale bar = 20 µM. **f** Morphological profile heat map of ochratoxin and the PIK3CA antagonist buparlisib and taselisib at highest tested concentration represented by normalised feature values (B-scores). B-scores were clipped to the range [-6, +6] for visualization. Channel, region and module are encoded by colour as indicated in panel b. Pearson correlation coefficient (r) between ochratoxin and the other two compounds are indicated. A black square highlights lysosome (lyso)-related features. **g** Representative images of compound-induced mitochondrial changes in RPTEC treated with omeprazole, diclofenac, and NPA. Mitochondria and DNA channel are displayed. Scale bar = 20 µM. Lyso=lysosomes; Mito=mitochondria; ER=endoplasmic reticulum; Golgi=Golgi apparatus.

This difference in exposure level may partly explain the absence of detectable phenotypic activity in our CPP assay. Further, as in the CPP assay only one time point was measured (after 24 h), it cannot be excluded that prolonged exposure time would induce observable changes.

To identify compound-specific morphological alterations and groups of compounds causing similar morphological responses, two approaches were employed. First, a morphological profile heatmap based on normalised feature values (B-scores) and hierarchical clustering using Pearson correlation was generated using maximum non-cytotoxic concentrations (Fig. 2b). Second, to integrate dose-dependent effects across all non-cytotoxic concentrations, benchmark concentrations (BMCs) were calculated for individual features as previously described for CPP (von Coburg et al. 2025; Wedler et al. 2026a). The BMC represents the lowest concentration at which a significant change in a specific feature is detected. For each compound and CPP channel, the proportion of features showing significant responses to a compound compared with all tested compounds was summarised and used to generate a BMC proportion map (Fig. 2c). Exemplary compounds with organelle-specific phenotypes are highlighted with coloured compound names in Fig. 2c, b and are described in more detail below.

Overall, most compounds affected several morphological features with varying effect sizes. In both plots, cadmium dichloride and potassium dichromate showed the strongest responses across all channels (Fig. 2b, c) consistent with their high percentage of active features as shown in Fig. 2a. Besides strong perturbations across many organelles (for example, cadmium dichloride and potassium dichromate), several more organelle-specific effects, namely on DNA, ER/lysosome and mitochondria, were observed and compared, forming the basis of the phenotypes described below.

#### DNA phenotypes

First, we examined compounds that induced pronounced DNA phenotypes: Both doxorubicin and indomethacin clustered together when comparing their proportion of significant BMCs in the different channels showing predominantly affected DNA features (compound name is highlighted in green in Fig. 2c). However, their morphological profiles clustered in separate groups (Fig. 2b), indicating differences in the cellular responses to DNA disruption. Doxorubicin elicited predominant DNA and RNA responses as shown in the morphological profile heatmap (Fig. 2b), and grouped together with gentamicin and diethyl maleate (Fig. 2b). This is consistent with known capacity of doxorubicin (Kciuk et al. 2023), gentamycin (El Mouedden et al. 2000; Elliott and Jiang 2019) and diethyl maleate (Cardozo-Pelaez et al. 2002; Geter et al. 2012) to induce DNA fragmentation and/or oxidative DNA stress. Indomethacin also displayed a distinct DNA phenotypic pattern. However, indomethacin showed increased DNA features whereas doxorubicin exhibited decreased DNA features (Fig. 2b). This dichotomy was confirmed by box plots and representative CPP images of DNA staining: indomethacin induced a concentration-dependent increase in nuclear DNA intensity, while doxorubicin caused a dose-dependent decrease (Fig. 2d). Doxorubicin damages DNA by intercalation (Nedelnikova et al. 2025) and TOP2 inhibition (Tewey et al. 1984) and generates oxidative stress (Panis et al. 2012; Kciuk et al. 2023), whereas, indomethacin, a non-steroidal anti-inflammatory drug, suppresses COX-1/2 but has also been reported to intercalate DNA (Husain et al. 2017). The divergent phenotypes therefore likely reflect different MoAs of these compounds, which however can be distinguished by the high granularity of the phenotypic profiling method.

Carboplatin and cisplatin grouped together (Fig. 2b, c) and are known to form covalent platinum-DNA adducts (Rabik and Dolan 2007), yet they elicited only modest DNA effects in the CPP assay along with notable lysosomal responses (Fig. 2b, c). This may reflect reports that DNA-damaging chemotherapeutics can induce lysosomal damage and disrupt lysosomal membranes (Leibbrandt et al. 1995; Toriyama et al. 2021; Yamashita et al. 2022; Tian et al. 2025) and that lysosomal sequestration and autophagy are evident for platinum compounds (Safaei et al. 2005; Rodrigo et al. 2019).

#### ER/lysosome phenotypes

The ER-stress inducers thapsigargin and tunicamycin (purple compound labels in Fig. 2b, c) showed similar phenotypic profiles and displayed high active BMC proportions not only in ER but also in lysosomal and mitochondrial features (Fig. 2c), consistent with literature linking the unfolded-protein response to ROS-mediated mitochondrial apoptosis (Guha et al. 2017) and to lysosomal transport inhibition (von Figura et al. 1979; Imort et al. 1983; Ganley et al. 2011).

Compounds with pronounced ER activity also included ochratoxin A, which showed a high proportion of active BMCs for ER and lysosomal features (Fig. 2c). To visualise compound potency in more detail, features were grouped into feature categories (FCs) (based on feature Channel, Region and Module) and the median BMC per category was calculated as described previously (von Coburg et al. 2025; Wedler et al. 2026a). Accumulation plots ranked categories by potency; ochratoxin A exhibited the lowest median BMCs for several lysosomal FCs followed by ER categories (Supplementary Fig. S3a), indicating strong potency for lysosomal effects. Correspondingly, ochratoxin-treated cells displayed increased lysosomal intensity and altered texture compared with DMSO solvent control in representative lysosome images (Fig. 2e).

Ochratoxin A activates PI3K/AKT and MAPK/ERK1-2 pathways and modulates autophagy (Akpinar et al. 2019; Izco et al. 2021). Given that the reference chemicals contain annotated PIK3CA antagonists, we compared their phenotypic profiles with that of ochratoxin A (Fig. 2f). High concordance, particularly for lysosomal features, was observed, with correlation score of 0.75 (ochratoxin A and taselisib) and 0.57 (ochratoxin A and buparlisib). This demonstrates that morphological profiling can link phenotypic signatures to a compound’s MoA.

#### Mitochondria phenotypes

In the BMC proportion map, several compounds, such as tunicamycin and ochratoxin A demonstrated above, showed combined high potency for mitochondrial effects with strong lysosomal responses. Puromycin also showed pronounced changes in mitochondrial features, but also in Golgi-related features, consistent with its reported effects on mitochondrial ATP synthesis (Price and Verner 1993) and Golgi complexes (Sturgess et al. 1975).

Compounds with strong mitochondria-specific activity included for example diclofenac, NPA and omeprazole (orange labeled compound names in Fig. 2b, c). Accumulation plots confirmed the lowest median BMCs for multiple mitochondrial feature categories for omeprazole and NPA (Supplementary Fig. S3b).

Although omeprazole grouped with diclofenac and NPA in the BMC proportion map (Fig. 2c), this compound had decreased responses in mitochondrial features, whereas NPA and diclofenac displayed increased responses in mitochondrial features (Fig. 2b). These trends were consistent across concentrations and were dose-dependent (Supplementary Fig. S3c). Representative mitochondrial channel images corroborated the quantitative data: omeprazole reduced mitochondrial staining intensity, while diclofenac and NPA increased it without affecting the DNA channel (Fig. 2g). Enhanced mitochondrial signal likely reflects early mitochondrial swelling and remodelling as previously described for these compounds (Masubuchi et al. 2002; Tatematsu et al. 2016; Jung et al. 2020), whereas the reduced signal with omeprazole aligns with loss of membrane potential and diminished dye uptake (Xiong et al. 2026).

In summary, the CPP assay enables fine-grained discrimination of morphological features, which can be linked to similar or opposing modes of modulation. Nephrotoxic compounds can be systematically grouped by the cellular compartment they affect, underscoring the utility of CPP to recapitulate organelle-specific phenotypes of nephrotoxic compounds.

### Morphological signatures of growth factors and their antagonists

Next, we examined the morphological profiles of the growth factors and cytokines tested in RPTEC CPP screen. Generally, the tested growth factors and cytokines elicited weaker morphological responses than the nephrotoxic compounds described above. BMP-4, FGF1, EGF and Wnt-3A induced only faint morphological changes. In contrast, activin A, TGF-β1, TNF-α and IL-1β caused noticeable feature changes (Fig. 3a, b). Among them, two prominent phenotypic patterns emerged. The first group comprised TGF-β1 and activin A, both members of the TGF-β superfamily. They produced the strongest alterations in actin-related features and yielded the highest correlation coefficient of 0.85 (Fig. 3a, c), which is consistent with their shared signalling pathways and known effects on actin remodelling (Boland et al. 1996; Nalluri et al. 2015). The second group included the cytokines IL-1β and TNF-α, which decreased features across all channels and showed a high correlation coefficient of 0.72 (Fig. 3a, c). Interestingly, TGF-β1 and activin A strongly increased actin-related features, whereas TNF-α and IL-1β decreased them (Fig. 3a). These opposite phenotypes likely stem from distinct MoAs: Activin A and TGF-β1 signal through the SMAD pathway (Morianos et al. 2019; Melchionna et al. 2021) and are implicated in regulated cytoskeletal reorganisation during processes such as epithelial-mesenchymal transition (Boland et al. 1996; Nalluri et al. 2015). In contrast, TNF-α and IL-1β are pro-inflammatory stressors that trigger actin remodelling via NF-κB signalling, promote formation of dense stress fibres, increase cellular stiffness, and disrupt junctional complexes (Campos et al. 2009; Kakiashvili et al. 2009; Lie et al. 2012; Chen et al. 2015). This example illustrates how divergent morphological signatures can be linked to underlying MoAs.

**Fig. 3.**
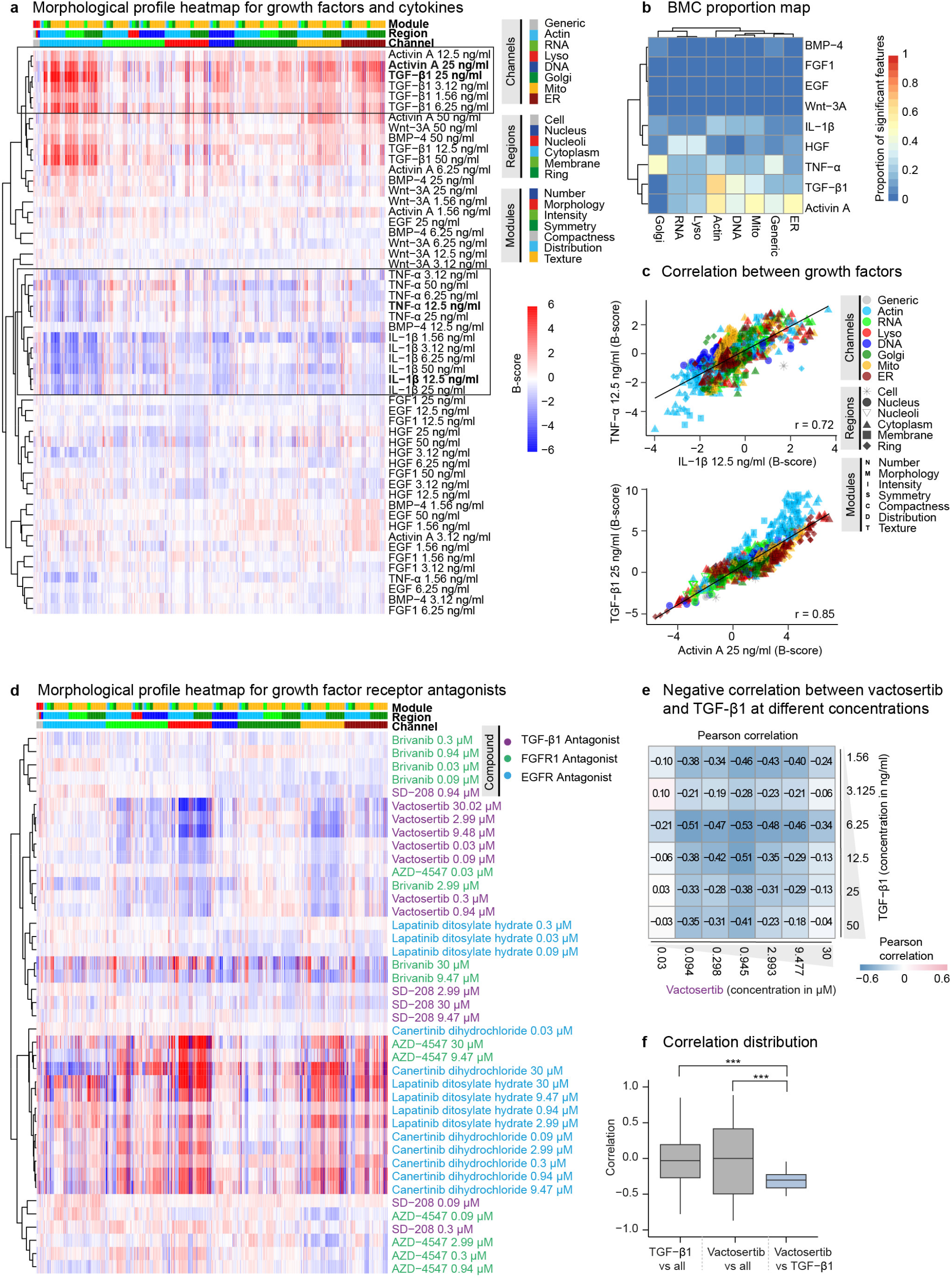
Morphological profiles of growth factors and their antagonists in RPTEC/TERT1. **a** Morphological profile heatmap representing normalised feature values (B-scores) of growth factors and cytokines at different tested concentrations. B-scores are clipped to the range [-6, +6] for visualisation. Modules, regions and channels are colour-coded. The dendrogram depicts Pearson correlation distance. Black squares indicate the groups of compounds specifically described in the main text. Bold text indicates concentrations shown in panel c. **b** BMC proportion map indicating proportion of single features that show a significant response compared to all other compounds (p<0.05) within each channel. Columns are hierarchically clustered using the Ward.D2 linkage method. **c** Correlation between morphological profiles of TNF-α and IL-β1 and between TGF-β1 and activin A. Symbols represent B-scores of single morphological features, with corresponding channel, region and module encoded by colour, shape and letter, respectively. r indicates Person correlation coefficient. Lyso=lysosomes; Mito=mitochondria; ER=endoplasmic reticulum; Golgi=Golgi apparatus. **d** Morphological profile heatmap representing B-scores of growth factor antagonists. Modules, regions and channels are colour-coded as in panel a. Dendrogram depicts Pearson correlation distance. **e** Pearson correlation between morphological profiles of TGF-β1 and the TGF-β1 antagonist vactosertib at different concentrations. Numbers inside the squares show Pearson correlation score, also indicated by colour; blue indicates negative and red positive correlation. **f** To assess whether TGF-β1 and vactosertib show a significant correlation,, Pearson correlation between their morphological profiles were calculated and compared with background correlations (each compound versus all others, excluding the pair) using a t-test. For this analysis all active concentrations were used. A concentration was defined as active when ≥1% of features have an absolute B-score ≥ 3. Asterisks (***) indicate a significant difference at p<0.001.

As the annotated reference chemicals compound set contains antagonists of TGF-β, EGFR and FGFR, we next compared their morphological profiles with those induced by the corresponding growth factors. We hypothesised that antagonists and agonists would exhibit systematic negative correlations.

First, we inspected the compound-specific heatmap summarising all concentrations of the six antagonists (Fig. 3d). Vactosertib produced a markedly stronger response than SD-208, reflecting its higher potency for TGF-β receptor ALK5 (IC₅₀ = 11 nM versus 48 nM) and its affinity for additional TGF-β receptor ALK4 (IC₅₀ ≈ 13 nM) (Uhl et al. 2004; Jin et al. 2014). Among the FGFR antagonists, AZD-4547 is highly selective (IC₅₀ ≤ 2,5 nM for FGFR1-3) (Gavine et al. 2012), whereas brivanib is a multi-kinase antagonist with primary activity against VEGFR2 (IC₅₀ = 25 nM) and weaker inhibition of FGFR1 (IC₅₀ = 148 nM) (Bhide et al. 2006). These potency differences were reflected in their distinct phenotypic signatures, resulting in their separation into different groups (Fig. 3d). For EGFR inhibition, canertinib is slightly more potent than lapatinib, although both have IC₅₀ values in the ∼10 nM range (Smaill et al. 2000; Rusnak et al. 2001). Consequently, they grouped closely (Fig. 3d) and showed only minor differences in morphological profiles.

Because EGF and FGF induced only faint morphological response (Fig. 3a, b), we selected only TGF-β1 for correlation with its antagonist. For the TGF-β1 antagonist, we chose vactosertib over SD-208 because it induced stronger morphological responses. We hypothesized that TGF-β1- and vactosertib-treated cells would have negatively correlated morphological profiles. Indeed, they showed a consistent but weak negative correlation across most tested concentrations (Fig. 3e). This correlation differed significantly from background correlations with all other chemicals (Fig. 3f). This example demonstrates that CPP captures the expected inverse relationship between the TGF-β1 receptor antagonists and agonists.

### RPTEC CPP profiling of target-annotated reference chemicals

To further explore the discriminatory power of CPP in RPTEC, we expanded the analysis to the 284 target-annotated reference chemicals. As many compounds target multiple proteins, the high-confidence target (with highest support level) for each compound was identified as previously described (Judson et al. 2019; Bundy et al. 2022), resulting in 137 targets in total (Fig. 4a). Since compound function depends on both the target protein and the mode of modulation, target identity and modulation (agonist or antagonist) were jointly defined as the compound MoA (Supplementary Table 1).

**Fig. 4.**
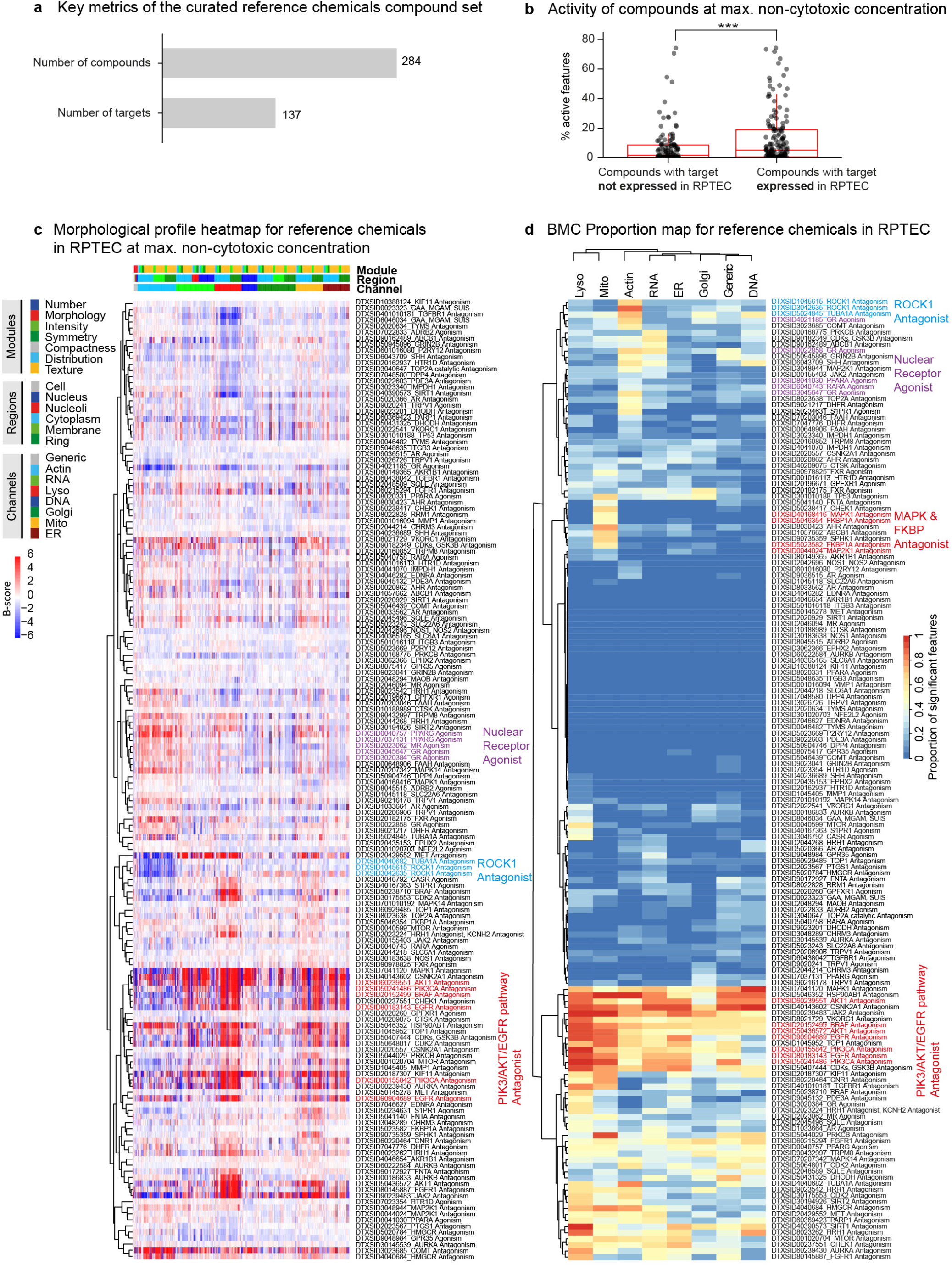
Screening of the annotated reference chemicals in RPTEC/TERT1. **a** Metrics of the compounds used in this screen: 284 compounds were annotated with 137 targets, each shared by at least two compounds. **b** The compounds were classified according to whether their target is expressed in RPTEC, based on mRNA expression data from the Human Protein Atlas. Bioactivity for each compound was assessed based on the percentage of active morphological features at the maximum non-cytotoxic concentration. Gray dots represent values for the individual tested compounds, median values (from n=5 biological replicates) are shown. Red box plots depict the median and first and third quartiles. Asterisks indicates significant difference based on Wilcoxon rank sum test with p ≤ 0.001. **c** Morphological profile heatmap of compounds at the maximum non-cytotoxic concentration using normalised feature values (B-scores). Only compounds with target expressed in RPTEC are depicted. Coloured compound labels highlight compounds mentioned in the main text. B-scores are clipped to the range [-6, +6] for visualization. **d** Proportion of active BMCs indicating the potency of compounds for organelles. Single-feature BMCs were derived from n = 5 biological replicates. Colour represents proportion of single features showing a significant response to the specific compound compared to all other compounds (p ≤ 0.05) within each channel. Compounds clustered by shared or functionally related protein targets are annotated by target class, with colour highlighting compound groups specifically mentioned in the main text. Only compounds with targets expressed in RPTEC are depicted. Lyso=lysosomes; Mito=mitochondria; ER=endoplasmic reticulum; Golgi=Golgi apparatus.

Because not all high-confidence targets are expressed in RPTECs according to the Protein Atlas mRNA data (https://www.proteinatlas.org) (Uhlen et al. 2015; Jin et al. 2023) (Supplementary Table 1), compounds were grouped based on whether their primary target was expressed in RPTECs. Those targeting expressed proteins induced significantly stronger morphological responses, affecting a greater number of features than compounds against targets that were not expressed (Fig. 4b). Nevertheless, several compounds whose primary target was absent still elicited substantial morphological activity, suggesting potential off-target effects or alternative MoAs (Fig. 4b).

In order to analyse the compounds whose targets are expressed in RPTEC in greater detail, we generated a morphological profile heatmap and a BMC proportion map as described above. Most of them exhibited weak morphological responses and low proportions of active BMCs (Fig. 4c, d). The largest group, containing many compounds with strong, multi-feature morphological effects, was characterised by pronounced lysosomal, mitochondrial and ER alterations and included kinase antagonists targeting EGFR, PIK3CA, and AKT1 (red compound labels in Fig. 4c). This group was also evident in the BMC proportion map, showing strong activity across all organelles with especially pronounced effects for lysosomes (red compound labels in Fig. 4d). In addition, two compound groups with opposing prominent actin phenotypes were observed in the morphological profile map. One group, characterised by increased actin-related features, consisted of compounds targeting nuclear receptors (NRs), including the glucocorticoid receptor (GR), mineralocorticoid receptor (MR), and peroxisome proliferator-activated receptor gamma (PPARγ) (purple compounds in Fig.4c). The other group, characterised by decreased actin-related features, comprised two ROCK1 antagonists and a TUBA1A (α-tubulin) antagonist (blue compounds in Fig. 4c). This finding is consistent with the established role of ROCK1 as a key regulator of cytoskeletal organisation (Amano et al. 2010) and of α-tubulin as a cytoskeletal protein and major component of microtubules (Janke 2014). These ROCK1 antagonists and NR regulators also grouped together in the BMC proportion map, exhibiting a high proportion of active actin BMCs (Fig. 4d). Furthermore, the two FKBP1A antagonists and the MAPK antagonists were characterised by pronounced mitochondrial effects in the BMC proportion map (Fig. 4d). The enrichment of mitochondrial phenotypes among MAPK- and FKBP1A antagonists is consistent with the known role of these signalling networks in regulating mitochondrial homeostasis. MAPK pathways, particularly ERK and p38, directly modulate mitochondrial function and stress responses (Sarg et al. 2024), while FKBP1A-associated mTOR signalling integrates mitochondrial and metabolic cues (Desai et al. 2002).

In summary, RPTEC exhibited pronounced responses to several classes of kinase antagonists. Compounds showed pathway-specific morphological signatures, particularly for the EGFR–PI3K/AKT signalling axis, while also capturing characteristic phenotypes associated with cytoskeletal and nuclear receptor regulation.

### Cell-type-specific morphological differences and CPP responses between RPTEC and U2OS cells

As the bone osteosarcoma cell line U2OS is the most frequently employed model for CP screens (Gustafsdottir et al. 2013; Bray et al. 2016; Nyffeler et al. 2020; Pahl et al. 2023), we applied the CPP workflow used for RPTEC to U2OS in order to uncover cell-type-specific characteristics. The 284 annotated reference chemicals were screened at a single concentration corresponding to one of the 7 concentrations used in the RPTEC CPP screen (Fig. 5a, a’). The concentrations selected for the U2OS CPP screen were based on the cytotoxicity data from a separate U2OS screen described in our companion manuscript (Wedler et al. 2026b, in preparation) and, for several compounds, were lower than the MaxConc used in the RPTEC CPP screen. Therefore, subsequent cross-cell-line comparisons were restricted to concentrations that were tested in both cell lines.

**Fig. 5.**
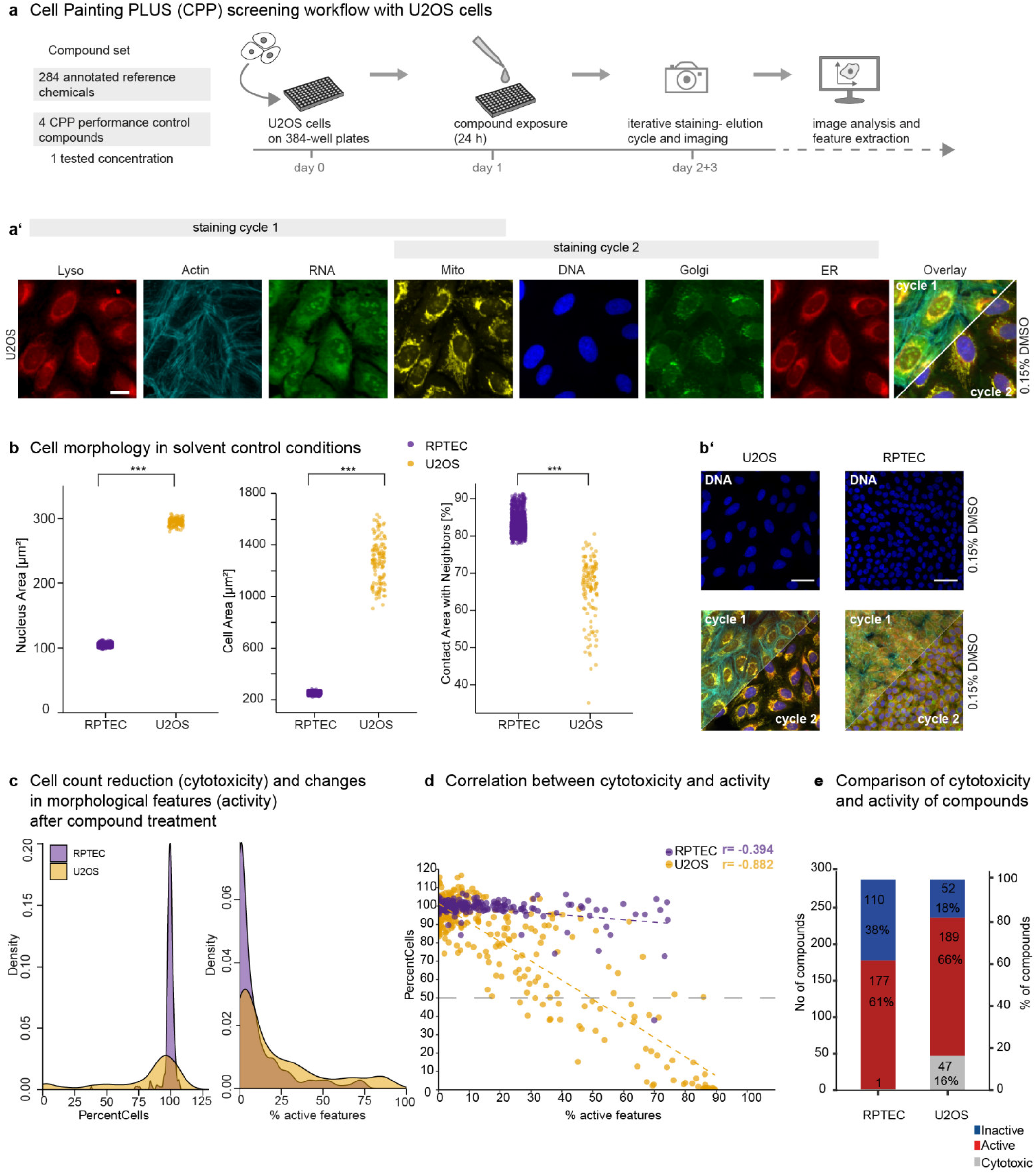
Cell-type-specific morphological differences between RPTEC/TERT1 and U2OS cells. **a** U2OS CPP screening workflow and representative images of solvent control (DMSO) in U2OS cells (a’). n = 4. Scale bar = 20 µm. Lyso=lysosomes; Mito=mitochondria; ER=endoplasmic reticulum; Golgi=Golgi apparatus. **b** Quantitative cell morphology comparison between U2OS and RPTEC in solvent control (DMSO). Nuclear and cell areas (in µm^2^) are shown. Dots represent the solvent control wells, pooled from all biological replicates. Asterisks indicate significant differences based on Wilcoxon rank-sum test with p ≤ 0.001. **b’** Representative images of RPTEC and U2OS cells in solvent control, depicting DNA channel (top) and overlay of the CPP staining cycle 1 and 2 (bottom). Note that a broader imaging area is displayed compared to panel a’ in order to provide an overview of cell density, scale bar = 500 µm. **c** Probability density of the percentage of total cells relative to solvent control (PercentCells) and of activity (expressed as % active features) after exposure to reference chemicals. A feature is considered active with B-score (RPTEC) or robust z-score (U2OS) ≥ 3. The y-axis indicates the relative probability density. Colours represent the different cell lines. **d** Scatter plots of PercentCells versus percentage of active features across RPTEC and U2OS cells. Correlation coefficients (r) are displayed. **e** Comparison of cytotoxicity and activity of reference chemicals across RPTEC and U2OS cells. A compound is considered cytotoxic if it results in reduced PercentCells below 50%. A compound is considered active if at least 1% of features shows a robust z-score or B-score ≥ 3 (representing activities ≥3xMAD of the DMSO control).

The U2OS screen proved robust and reproducible, as indicated by percent replicating score of 0.68 (Supplementary Fig. S4a) and absence of pronounced plate effects (Supplementary Fig. S4b). A subpopulation of cells, exhibiting markedly reduced nuclei size, indicative of dead or fragmented nuclei, was identified and excluded prior to robust z-score normalisation (Supplementary Fig. S4c).

Consistent with representative CPP images, densely packed, differentiated RPTEC exhibited significantly smaller nucleus and cell size, as well as a larger cell-to-cell contact area with neighbouring cells compared with confluent U2OS cells (Fig. 5b, b’).

Comparative analysis of compound-induced responses revealed that RPTEC experienced only modest reductions in the percentage of total cells relative to the DMSO control; the majority of compounds maintained ∼100 % total cells and exhibited 0-25% active features (Fig. 5c). In contrast, more compounds in U2OS cells reduced cell count and showed an increase in the number of active features. Moreover, in U2OS the cell count was strongly negatively correlated with the number of active features (r = −0.88), whereas this relationship was weaker in RPTEC (r = −0.39) (Fig. 5d). When the same concentrations were considered, 47 of the tested compounds were cytotoxic in U2OS, whereas only one compound met this criterion in RPTEC. U2OS also displayed a slightly higher proportion of active and non-cytotoxic compounds compared with RPTEC (66 % vs 61 %) (Fig. 5e). This supports the idea that RPTEC is more resilient to the chemical treatment than U2OS.

The increased resilience of RPTEC relative to U2OS, HepG2, and MCF-7 cells has been consistently observed in a previous CPP screen employing performance-control compounds (von Coburg et al. 2025), suggesting an intrinsic cellular characteristic of RPTEC.

Because RPTEC are considerably smaller than U2OS cells, we investigated whether their diminished morphological response reflects a genuine biological effect or merely arises from detection limits imposed by small cell size. To this end, we compared RPTEC morphology with that of the densely seeded adenocarcinoma epithelial line MCF-7 and the subconfluent U2OS cells, which was included in a previously published CPP screen comprising 15 performance control compounds (Supplementary Fig. S5a) (von Coburg et al. 2025). RPTEC and MCF-7 cells possess comparable nuclear and cellular dimensions, both markedly smaller than those of U2OS (Supplementary Fig. S5b). Nevertheless, MCF-7 cells exhibited a higher number of highly active compounds and a strong negative correlation between cytotoxicity and activity (r = −0.94) compared with RPTEC (Supplementary Fig. S5c, d). These findings indicate that the reduced response and heightened resistance observed in RPTEC are not attributable to imaging or detection constraints, but rather represent a true biological phenomenon.

In summary, RPTEC demonstrate a markedly lower propensity for morphological perturbation and a higher intrinsic resistance to cytotoxicity than U2OS. This resistance cannot solely be explained by the reduced cell size, as a comparably small, densely seeded cell line (MCF-7) exhibits pronounced morphological responses. Consequently, the attenuated phenotype of RPTEC reflects genuine cell-type-specific biology rather than methodological artefact, underscoring the importance of incorporating physiologically relevant renal epithelial models in toxicological screening.

### Cell-type-dependent phenotypic response to compounds with distinct MoAs

Given the differences in cell morphology and compound responses between RPTEC and U2OS cells, we asked whether the response to compounds with distinct MoAs also differed between the two cell types. Pearson correlations were calculated for compounds that share the same annotated MoA and are present in both datasets (Fig. 6a). For this approach, multi-annotated targets of the compounds (column “MoA_complete” in Table S1) were retained to account for polypharmacology. High correlation scores between compounds with the same MoA indicate high concordance between morphological profiles of compounds targeting the same molecular target and thus good MoA matching, whereas low correlation suggests dissimilar morphological profiles. The overall correlation between the two cell lines was low (r = 0.17), indicating substantial differences in MoA concordance (Fig. 6a). The only target that showed a high correlation in both cell types (r > 0.75) was IMPDH1 (highlighted in red in Fig 6a), a key enzyme in DNA synthesis, signal transduction, energy metabolism, and glycoprotein biosynthesis (Wang and Hedstrom 1997). RPTEC exhibited higher correlations for compounds targeting PIK3CA (highlighted in purple) and ROCK1 and SQLE compared with U2OS (Fig. 6a), consistent with the phenotypes shown in Fig. 4c, d. Conversely, U2OS cells more accurately matched antagonists of TOP1/2A, (highlighted in blue in Fig. 6a) and modestly higher correlations for mTOR, and DPP4.

**Fig. 6.**
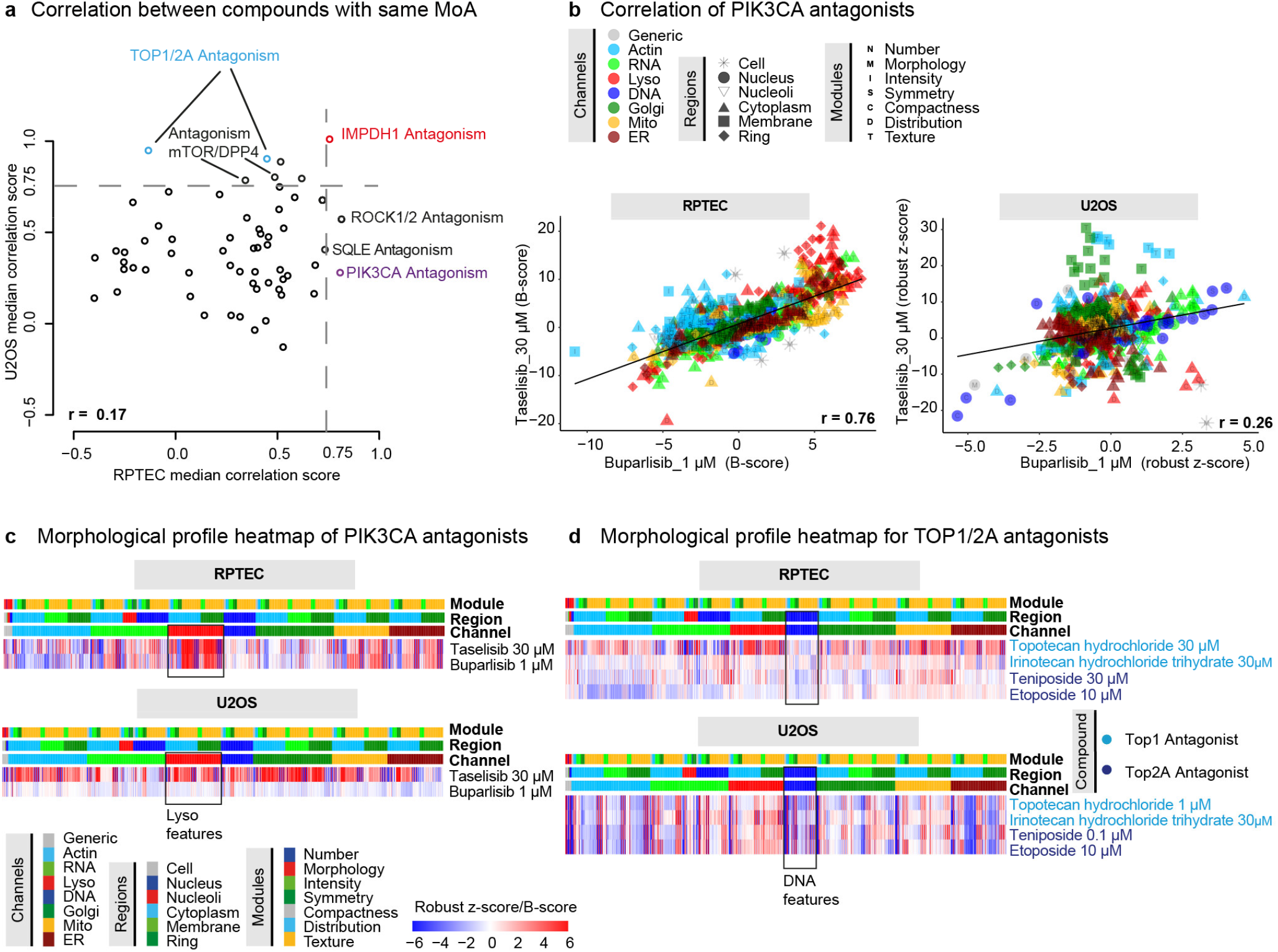
Cell-type-specific phenotypic response to compounds with distinct MoAs in RPTEC/TERT1 and U2OS. **a** Pearson correlation scatter plots for compounds sharing the same MoA, comparing U2OS and RPTEC data. Pearson correlation was calculated between morphological profiles of compounds sharing the same MoA. Each dot represents the correlation score for an individual MoA, comparing the U2OS data (y-axis) with the RPTEC data (x-axis). Dotted lines mark the threshold of 0.75, indicating high correlation. Coloured labels indicate the MoAs specifically discussed in the main text. **b** Correlation between PIK3CA antagonists in RPTEC and U2OS cells. Symbols represent normalised values of single morphological features, corresponding channels, regions and modules are encoded by colour, shape and letter, respectively. r indicates Person correlation coefficient. **c** Morphological profile heatmap (B-scores and robust z-scores) for PIK3CA antagonists in RPTEC and U2OS. Modules, regions, and channels are colour-coded. A black-outlined box highlights lysosome-related features. (B-scores and z-scores are clipped to the range [-6, +6], values > 6 are set to 6 and values < −6 are set to −6 for display). **d** Morphological profile heatmap for TOP1/2 antagonists in RPTEC and U2OS. Modules, regions, and channels are colour-coded as indicated in panel c. The black-outlined box highlights DNA-related features. Lyso=lysosomes; Mito=mitochondria; ER=endoplasmic reticulum; Golgi=Golgi apparatus.

As RPTEC showed higher correlation for kinase antagonists, particularly for PI3K signalling, we next focused on compounds from this class. The correlation between the PIK3CA antagonists taselisib and buparlisib was high (r = 0.76) in RPTEC, but lower (r = 0.26) in U2OS at the same tested concentrations (Fig. 6b). RPTEC tolerated a higher maximum non-cytotoxic concentration of buparlisib (30 µM vs 1 µM in U2OS); using this higher concentration increased the correlation to r = 0.82 in RPTEC (Supplementary Fig. S6). Both PIK3CA antagonists elicited a strong morphological response in RPTEC, especially affecting lysosome-related features (Fig. 6c), consistent with literature linking PI3Kα signalling to lysosome modulation (Endicott et al. 2020; Jia et al. 2022). In U2OS, buparlisib induced fewer changes than taselisib, and both compounds showed weaker lysosomal alterations compared with RPTEC. In contrast, topoisomerase antagonists, which were better correlated in U2OS (Fig. 6a), produced pronounced morphological changes in U2OS, and high similarity in DNA-related features (Fig. 6d). RPTEC displayed only modest responses, particularly for DNA features, even when higher concentrations of topotecan and teniposide were applied. This aligns with the known reduced sensitivity of post-mitotic, differentiated cells to agents that inhibit DNA replication or cell proliferation (Grandela et al. 2007; Secreto et al. 2017). While U2OS cells remain in proliferative state, RPTEC are not after 10 days of differentiation.

In summary, both RPTEC and U2OS cell lines reliably matched morphological profiles for core-essential targets, such as IMPDH1. RPTEC demonstrated higher morphological similarity for kinase antagonists, particularly PIK3CA-directed agents, whereas U2OS exhibited higher activity and higher correlation for topoisomerase antagonists. These cell-type-specific differences underscore the need to select physiologically relevant models for accurate toxicological screening and MoA elucidation.

## Discussion

Although CP has become a widely adopted morphological profiling assay, its application for nephrotoxicity prediction remains limited. Most image-based nephrotoxicity studies in renal cells rely on conventional high-content imaging or targeted multiplex assays rather than the canonical CP protocol (Wu et al. 2009; Li et al. 2013; Li et al. 2017; Jochum et al. 2025). As human proximal tubule (hPT) epithelial cells are a principal site of drug-induced nephrotoxicity, PT-derived cell lines such as HK-2, LLC-PK1, and RPTEC are widely used in single- or combined-endpoint assays, including cell viability (MTT, resazurin, LDH), apoptosis (caspase-3/7), cytokine expression, marker protein analysis and mitochondrial membrane potential (MMP) (Li et al. 2013; Li et al. 2017; Jochum et al. 2025). Beyond these targeted functional endpoints, transcriptomic approaches have emerged as powerful high-throughput tools for detecting early molecular perturbations and elucidating mechanisms of nephrotoxicity (van Kessel et al. 2025).

In the present work, we applied high-content morphological profiling using Cell Painting PLUS (CPP) with RPTEC, enabling fine-grained characterisation of compound-specific phenotypic effects. To this end, we evaluated the activity and similarity (correlation) of a set of nephrotoxic compounds, cytokines and growth factors together with a selected annotated reference chemicals compound set covering diverse signalling pathways and molecular functions.

Comparison with prior studies in immortalized proximal tubule cells (HK-2) and pseudo-immortalised primary proximal tubule epithelial cells (SA7K) highlights differences attributable to both cell model and assay endpoints. Previous reports assessed compound effects primarily via cytotoxicity or viability assays rather than multiplex phenotypic profiling (Wu et al. 2009; Li et al. 2017). In the present RPTEC-CPP screen, cadmium dichloride induced strong multiparametric alterations already at 1.25 µM, whereas in published studies using SA7K cells showed only weak viability impairment (IC₅₀ ≈ 65 µM) (Li et al. 2017). Similarly, gentamicin, which showed no cytotoxicity up to 1 mM in HK-2 cells (Wu et al. 2009), induced significant DNA, RNA, and lysosomal changes in our RPTEC-CPP screen at 300 µM. These observations are consistent with the capacity of CPP to detect sub-cytotoxic phenotypic perturbations that viability endpoints do not capture; however, direct comparison of intrinsic assay sensitivity is limited because RPTEC, HK-2, and SA7K differ in differentiation state and transporter expression. This cell-model dependence is further illustrated by mitochondrial responses to tetracycline and puromycin, which elicited marked effects in SA7K but were absent in HK-2 (Li et al. 2017). Consistent with this, puromycin and tetracycline also induced mitochondria-related morphological changes in the RPTEC-CPP screen presented here. Compared with SA7K, HK-2 cells exhibit absent or markedly reduced expression of several key renal transporters, which likely contributes to their lower responsiveness and predictive performance in nephrotoxicity assessment (Li et al. 2017). The differentiated RPTEC express transporter transcripts at levels comparable to primary human proximal tubule cells and showed reduced sensitivity to cyclosporin A (Aschauer et al. 2015). This is consistent with only mild phenotypic changes upon cyclosporin A treatment observed in the CPP screen presented here (Fig. 2a–c). Thus, RPTEC and SA7K both better represent core proximal tubule functions, such as renal transport, than HK-2, underscoring the importance of selecting physiologically relevant cell models. Together, these findings indicate that the differences observed across studies reflect not only the ability of CPP to reveal compound-induced alterations below cytotoxic thresholds, but also intrinsic variations in cellular susceptibility and functional maturity between proximal tubule models.

The published parallel transcriptomic profiling of the same nephrotoxic set in RPTEC (van Kessel et al. 2025) allows direct comparison of MoA signatures between gene expression and the CPP data shown here. Consistent MoA markers including DNA damage (doxorubicin, gentamicin), mitochondrial dysfunction (omeprazole), and ER stress (tunicamycin) were identified by both approaches, arguing that both methods yield concordant biological signatures. While transcriptomic analysis identified a core set of transcription factors perturbed by the majority of nephrotoxicants, providing mechanistic insight into the observed cytotoxicity (van Kessel et al. 2025), CPP further resolved phenotypic differences between compounds with related or divergent mechanisms. For example, doxorubicin and indomethacin both affected DNA-related channels but in opposite directions, whereas omeprazole and diclofenac produced distinct mitochondrial signatures. Thus, transcriptomics explained why the cells responded, whereas CPP revealed how those responses manifested morphologically.

CP studies have used U2OS cells because their large, well-spread cytoplasm and nucleus facilitates image segmentation, feature extraction, and morphological profiling. A recent comparison of U2OS and HepG2 cells reported that U2OS exhibited higher activity and greater sensitivity to compound-induced cytotoxicity (Wolff et al. 2025). However, sensitivity alone is not sufficient when selecting a cell model for MoA prediction. The ability to accurately group compounds with shared mechanisms, as reflected by biosimilarity, is an equally important performance criterion. A comparative study across six cell lines using CP assays demonstrated that greater phenotypic activity in a cell line did not necessarily translate to stronger morphological similarity among compounds with the same MoA, or to more accurate prediction of their MoA (Heinrich et al. 2023).

In our previous study comparing U2OS, MCF-7, RPTEC, and HepG2 cells using 15 CPP performance control compounds, RPTEC exhibited greater resilience to compound treatment than the other cell lines (von Coburg et al. 2025). This lower cytotoxicity and reduced activity relative to U2OS were confirmed in the present study using the reference chemicals compound set. Because RPTEC and MCF-7 cells were cultured at high density to maintain physiological morphology, whereas U2OS cells displayed a larger cell size and more sparsely distributed cell monolayer, we investigated whether differences in cell size and morphology could explain the lower sensitivity of RPTEC. Interestingly, although MCF-7 cells exhibited a morphology more similar to RPTEC than to U2OS, being smaller and grown in a more confluent cell layer, they showed higher activity and cytotoxicity than RPTEC, with a strong correlation between these two metrics comparable to that observed in U2OS cells. These findings indicate that the reduced sensitivity of RPTEC cannot be explained simply by their compact morphology or potential limitations in image segmentation and feature extraction. A more plausible explanation is the more physiological expression of xenobiotic transporters in RPTEC (Aschauer et al. 2015; Saib et al. 2021; Caetano-Pinto et al. 2022). The balance between uptake and efflux transporter activity determines intracellular drug exposure and, consequently, susceptibility to compound-induced toxicity. For some compounds, active efflux may reduce intracellular exposure, thereby attenuating cytotoxicity. For others, active uptake may accumulate them in the cells and enhance cytotoxicity.

RPTEC showed strong predictive performance for kinase antagonist activity, especially antagonists targeting the PI3K pathway components showed similar phenotypes and higher Pearson correlations than in U2OS. These findings are consistent with our accompanying studies with transcriptomic analysis (van Kessel et al. 2026), suggesting that RPTEC, as a primary-derived kidney cell line, may be biologically more responsive to kinase pathway perturbation. Conversely, U2OS showed better concordance for DNA-replication antagonists compared to RPTEC. This aligns with the post-mitotic nature of RPTEC, which are potentially less sensitive to DNA-replication targets.

Overall, our results demonstrate that RPTEC-based CPP combines physiologically relevant cell biology with high-dimensional phenotypic profiling to enable sensitive detection and characterisation of nephrotoxic compounds. Although RPTEC exhibited lower overall cytotoxicity and morphological activity than U2OS cells, their kidney-specific phenotype makes them a more biologically relevant model for studying nephrotoxicity.

Beyond conventional toxicity endpoints, CPP distinguished subtle phenotypic differences between compounds with related or opposing mechanisms of action and complemented transcriptomic analyses by linking molecular perturbations to cellular phenotypes. This integrated approach provides a promising platform for mechanism-informed nephrotoxicity screening and may contribute to improved preclinical safety assessment.

## Acknowledgments

We acknowledge our colleagues from BfR/Bf3R for their scientific input and valuable comments on the manuscript.

## Data availability

Data will be made publicly available upon publication in a scientific journal.

## Supporting information

**Supplementary Table 1: Annotated reference chemicals with curated and complete mechanisms of action (MoAs).**

Columns list the target, the type of modulation (Agonist = A, Antagonist = AT; where literature data support both activities, the entry is shown as A/AT), compound identity confirmation by mass spectrometry (MS-Identified) as described in van Kessel et al (2026) and the expression status of the corresponding mRNA in RPTE (mRNA_Expressed_RPTEC). mRNA expression data is retrieved from the Human Protein Atlas (https://www.proteinatlas.org/humanproteome/cell+line/data#cell_lines). A gene is considered not expressed when reported as zero TPM (transcripts per million). Note that parts of this table are also included in our companion manuscript (Wedler et al., 2026b in preparation) and are reproduced here for completeness.

**Supplementary Fig. S1.**
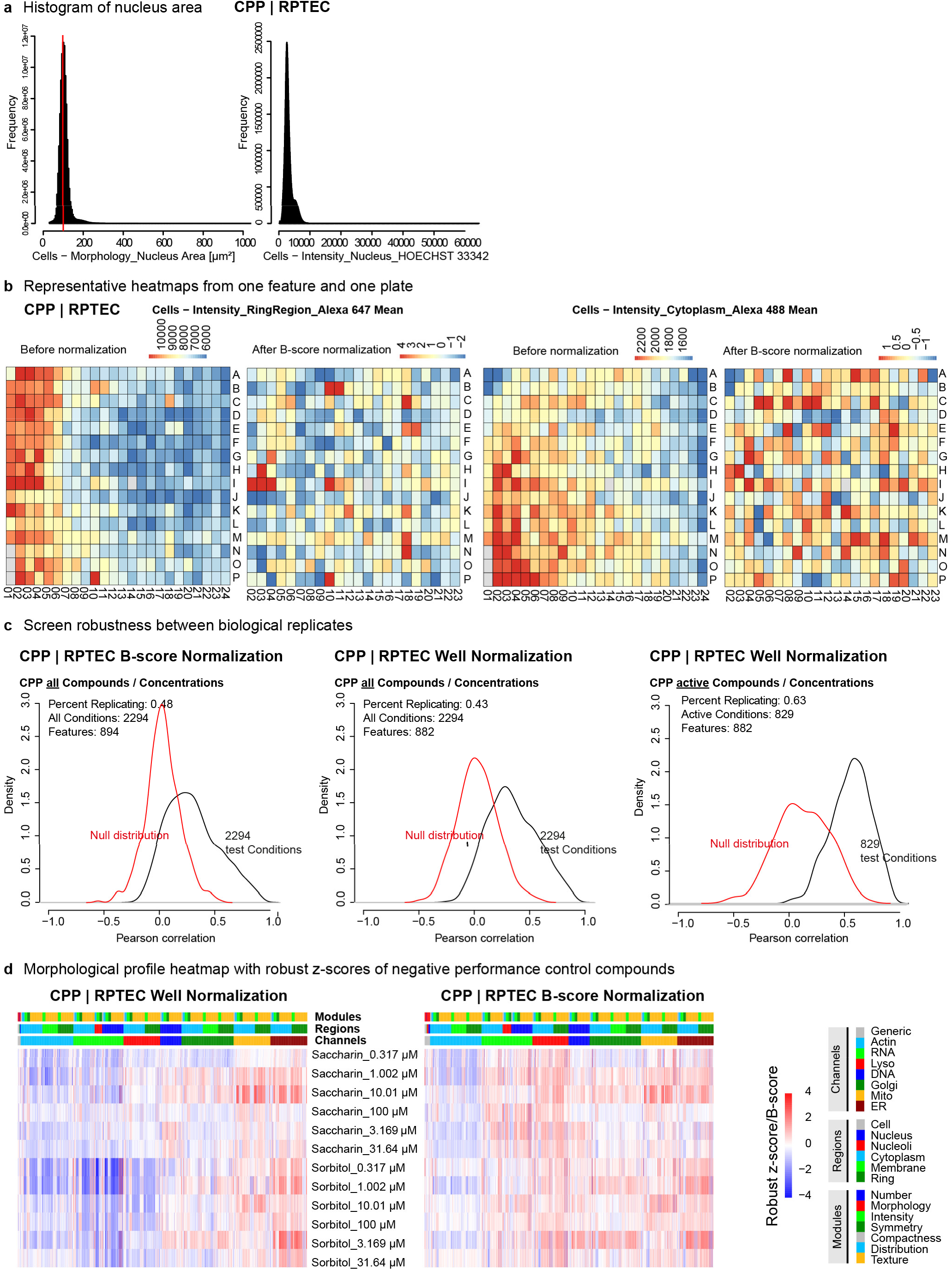
Evaluation of the RPTEC/TERT1 CPP screen. **a** Histograms of nuclear area and DNA signal intensity in nuclei across all biological replicates of the RPTEC/TERT1 CPP screen at single-cell level. **b** Representative heatmap visualising the mean signal intensity per well for the Alexa 647 (lysosome stain) channel on one representative screening plate, shown before normalisation and after B-score normalization. **c** Percent replicating after B-score and well-level normalisation for all conditions and active conditions only. **d** Morphological profile heatmap after well-level normalisation (robust z-score) or B-score normalisation (B-score) for CPP performance negative control compounds (sorbitol and saccharin), with respective concentrations (µM) indicated. Channels, regions, and modules are colour-coded. (B-score/z-score are clipped to the range [-4, +4] for display). Lyso=lysosomes; Mito=mitochondria; ER=endoplasmic reticulum; Golgi=Golgi apparatus.

**Supplementary Fig. S2.**
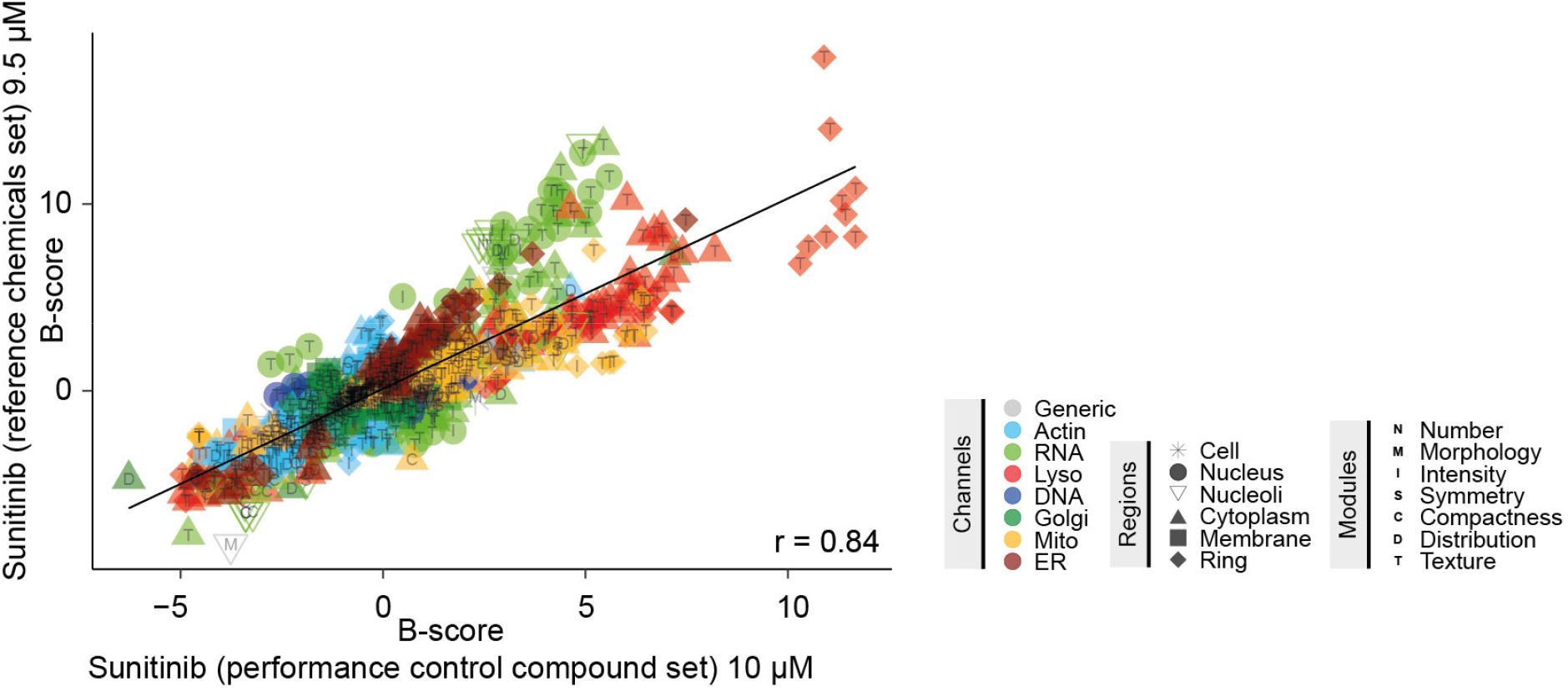
Correlation between sunitinib from different compound sets indicate screen robustness in RPTEC/TERT1. Sunitinib was included once in the reference chemicals compound set and once in the performance control compound set. Correlation was calculated between the morphological profiles of the compound. Symbols represent normalised feature values (B-scores) for single morphological features. Corresponding channels, regions, and modules are encoded by colour, shape, and letter, respectively. “r” indicates Person correlation coefficient. Lyso=lysosomes; Mito=mitochondria; ER=endoplasmic reticulum; Golgi=Golgi apparatus.

**Supplementary Fig. S3.**
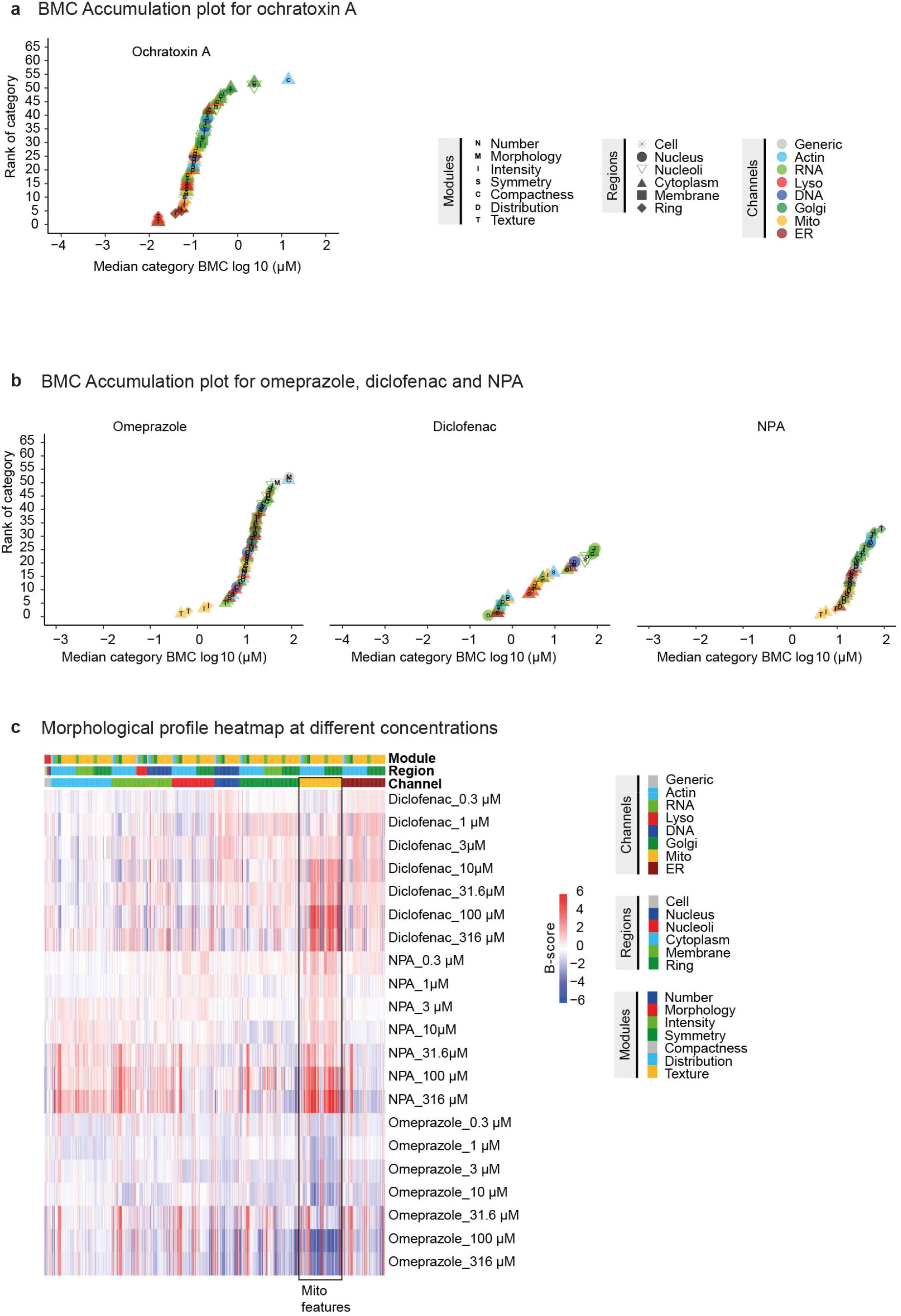
Feature category BMC and morphological profiles of omeprazole, diclofenac, NPA and ochratoxin A. **a** BMC accumulation plot for ochratoxin A: Features were grouped into feature categories according to channels, regions, and modules. Each data point represents the median BMC for one feature category in a concentration-dependent sequence (rank). Only feature categories including at least 30% features with active BMC are visualised in the plots. **b** BMC accumulation plot for omeprazole, diclofenac and NPA, as described in panel a. **c** Morphological profile heatmap representing single normalised features (B-scores) for diclofenac, NPA and omeprazole at different tested concentrations. Modules, regions and channels of the respective features are colour-coded. (B-scores are clipped to the range [-6, +6], values > 6 are set to 6 and values < −6 are set to −6 for display). Lyso=lysosomes; Mito=mitochondria; ER=endoplasmic reticulum; Golgi=Golgi apparatus.

**Supplementary Fig. S4.**
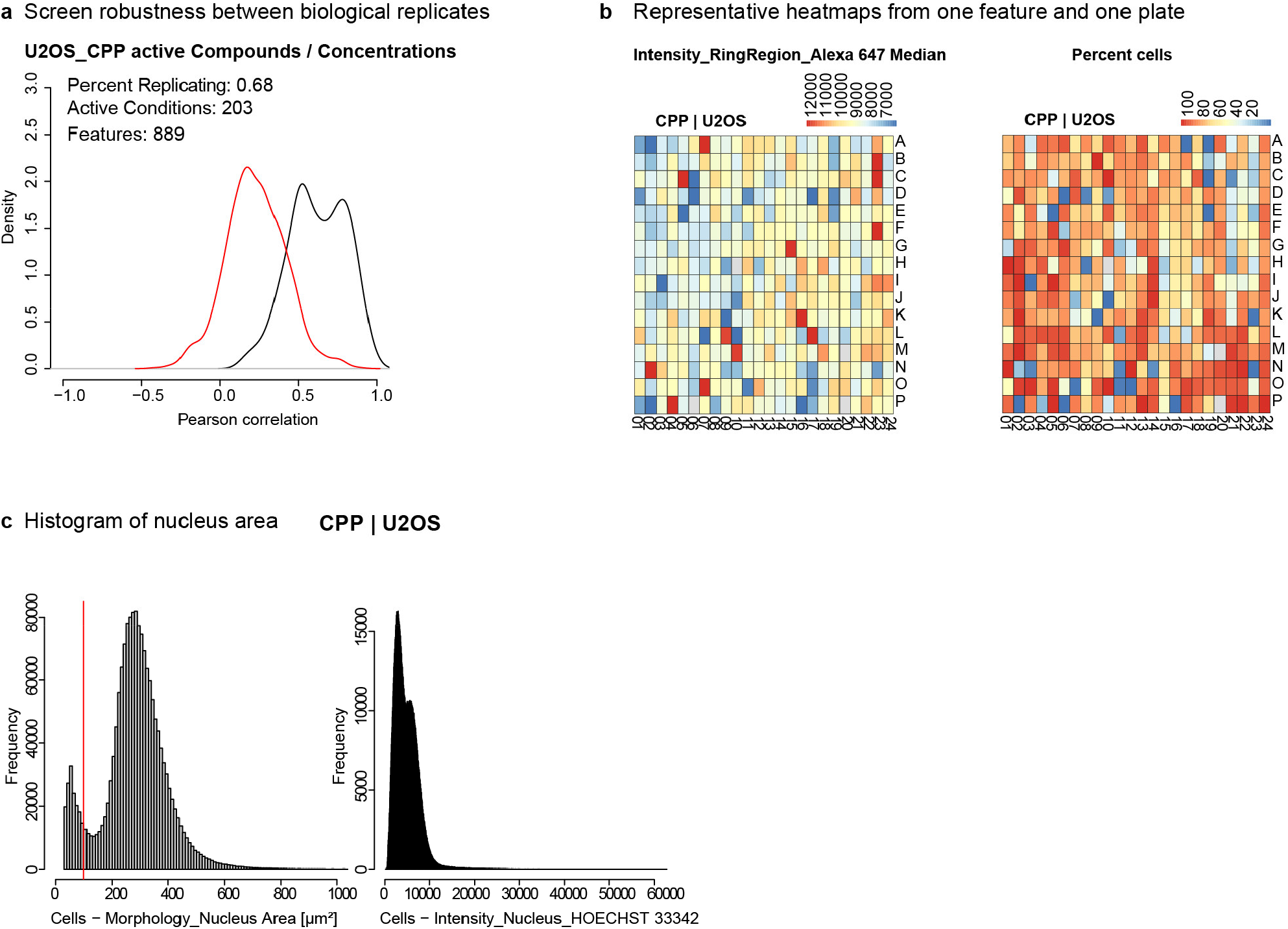
Evaluation of U2OS CPP screen. The plots were originally included in our companion manuscript (Wedler et al., 2026b) and are reproduced here for completeness. **a** Percent replicating score for active compounds at non-toxic concentrations across four biological replicates, demonstrating screening robustness under active conditions. **b** Representative heatmap showing the mean signal intensity per well in RingRegion for the channel Alexa 647 (lysosome stain) and the percentage of total cell numbers normalised to DMSO control on one screening plate for U2OS CPP. Unnormalised raw values for the lysosome stain feature are displayed. **c** Histograms of nuclear area and DNA signal (HOECHST) intensity. For U2OS cells, nuclei sizes were filtered using a nuclear area range of 120-1000 µm^2^ to exclude dead cells or cells with fragmented nuclei.

**Supplementary Fig. S5.**
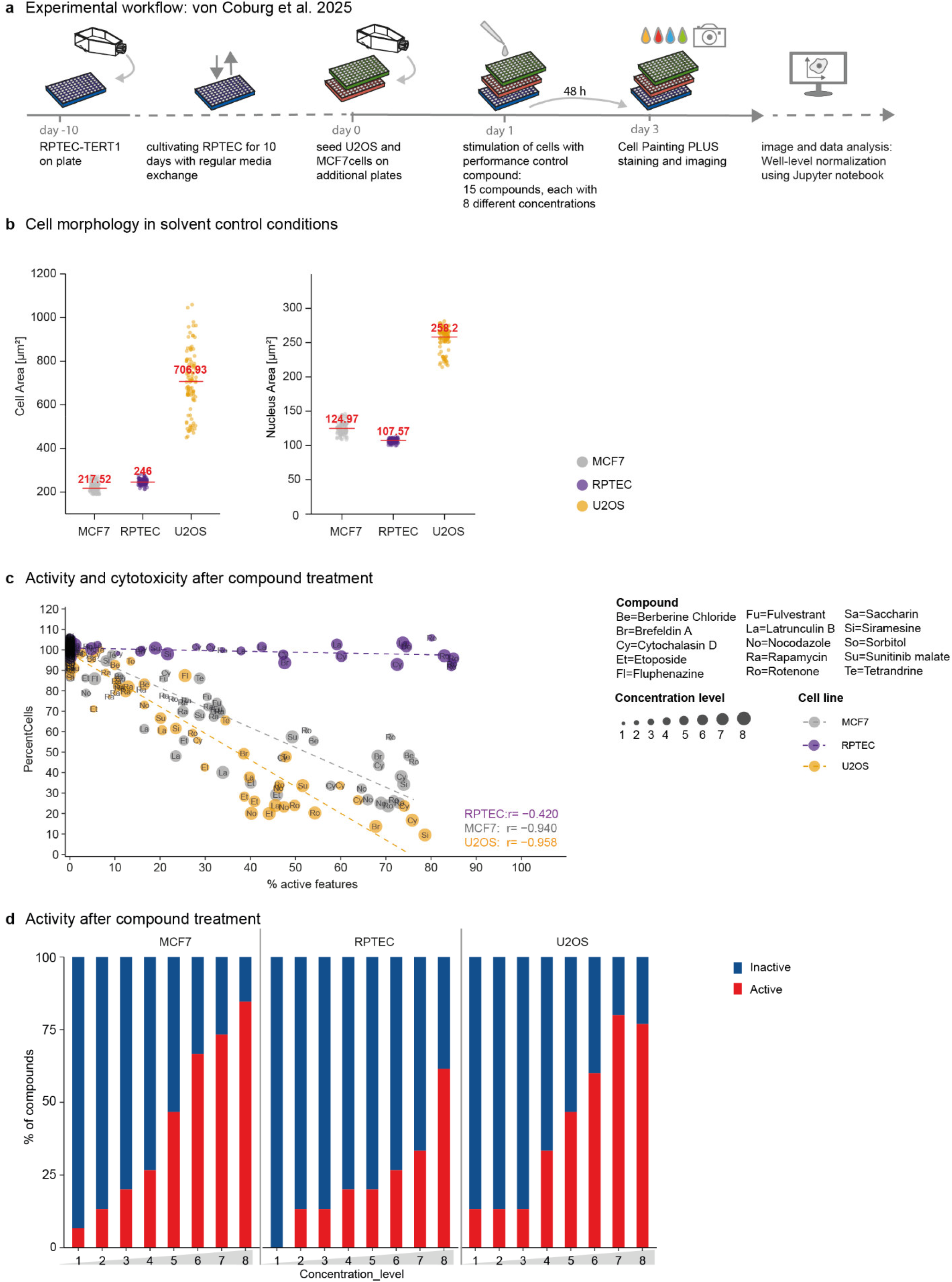
Comparison of cell morphology and activity between MCF7, RPTEC/TERT1 and U2OS cells. The data shown in this figure were initially published in von Coburg et al. (2025) and re-analysed here for the purpose of direct cell-line comparison. **a** Workflow of the Cell Painting PLUS (CPP) screen published in von Coburg et al., 2025: RPTEC/TERT1, MCF7/vBos and U2OS cells were cultivated on multi-well plates and exposed to a set of 15 control compounds for 48 h. Plate layout and compound concentrations are described in detail in the original publication. Tested compounds are listed in panel c. The data were normalised as described in the present publication using the Jupyter notebook. **b** Cell and nuclear areas in µm^2^ in solvent control (DMSO). Individual points represent median value per single well, showing all wells pooled from all replicates (4 independent biological replicates, each with 24 DMSO wells). Red lines and red numbers indicate median. Cell lines are colour-coded. **c** The percentage of total cells relative to solvent control condition (PercentCells) and percentage of active features after exposure to the 15 CPP performance control compounds at all tested concentrations. A feature is considered active with a robust z-score ≥ 3 (representing activities ≥ 3×MAD of the DMSO control). Single points indicate the median value for each tested condition from all replicates (4 independent biological replicates, each with 3 technical replicates). Compound names are encoded by letters, concentrations are encoded by dot size. Cell lines are colour-coded. “r” indicates Pearson correlation coefficient. **d** Activity of compounds for the different cell lines: Proportion of active compounds across all tested concentrations (level 1 = lowest tested concentration, level 7 = highest tested concentration). A compound is considered active when at least 1 % of all features have a robust z-score ≥ 3 (representing activities ≥ 3×MAD of the DMSO control).

**Supplementary Fig. S6.**
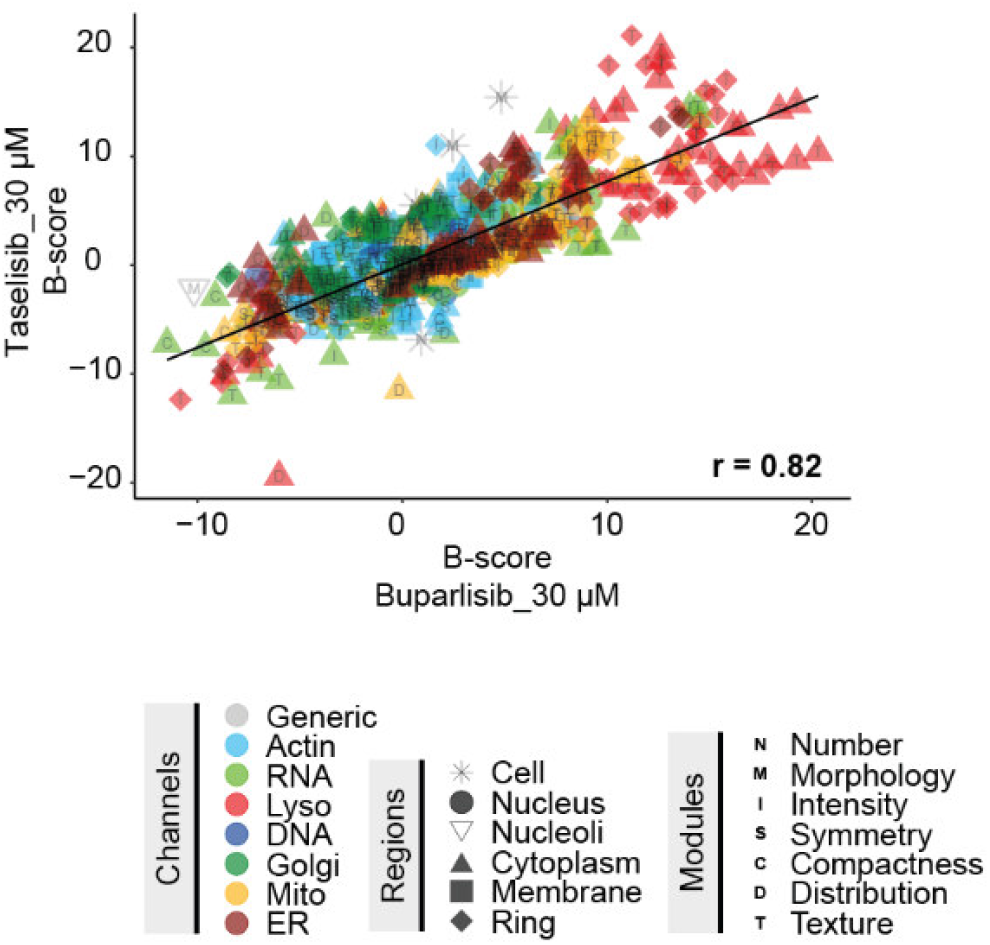
Correlation of PIK3CA antagonists in RPTEC/TERT1. Correlation between taselisib (30 µM) buparlisib (30 µM) and in RPTEC. Symbols represent normalised feature values (B-scores) of single morphological features. Corresponding channels, regions, and modules for each feature are encoded by colour, shape, and letter, respectively. ‘r=0.82’ indicates Pearson correlation coefficient. Lyso=lysosomes; Mito=mitochondria; ER=endoplasmic reticulum; Golgi=Golgi apparatus.

